# COCOA: Coordinate covariation analysis of epigenetic heterogeneity

**DOI:** 10.1101/2020.07.09.195289

**Authors:** John T. Lawson, Jason P. Smith, Stefan Bekiranov, Francine E. Garrett-Bakelman, Nathan C. Sheffield

## Abstract

A key challenge in epigenetics is to determine the biological significance of epigenetic variation among individuals. Here, we present Coordinate Covariation Analysis (COCOA), a computational framework that uses covariation of epigenetic signals across individuals and a database of region sets to annotate epigenetic heterogeneity. COCOA is the first such tool for DNA methylation data and can also analyze any epigenetic signal with genomic coordinates. We demonstrate COCOA’s utility by analyzing DNA methylation, ATAC-seq, and multi-omic data in supervised and unsupervised analyses, showing that COCOA provides new understanding of inter-sample epigenetic variation. COCOA is available as a Bioconductor R package (http://bioconductor.org/packages/COCOA).

## Introduction

Epigenetic data is inherently high-dimensional and often difficult to interpret. Because of the high dimensionality, it is common to group individual genomic loci into collections that share a functional annotation, such as binding of a particular transcription factor^[1–3]^. These genomic locus collections, or region sets, are analogous to the more common gene sets, but relax the constraint that data must be gene-centric. While gene set approaches may be applied to epigenetic data by linking regions to nearby genes^[4]^, this linking process is ambiguous and loses information because a regulatory locus may affect the expression of multiple genes or more distant genes. Alternatively, a region-centric approach is often more appropriate for epigenetic data, and there are now many region-based databases and analytical approaches^[1,2, 5–7]^, such as using region set databases for enrichment analysis^[1,7,8]^ or to aggregate epigenetic signals from individual samples across regions to assign scores of regulatory activity to individual samples or single cells^[2,3,6,9]^.

Region-based methods have provided complementary ways to annotate and understand epigenomic data, but they suffer from three drawbacks: First, it is common to ignore covariation between the epigenetic signal and continuous patient phenotypes, relying instead on differential signals between discrete sample groups. This approach loses information about the differences among samples within a group. Second, the use of discrete cutoffs for identifying significant epigenetic differences between samples loses information about the strength of covariation between epigenetic features and sample phenotype. Third, existing approaches are generally specific to certain scenarios (e.g. unsupervised analysis) or data types (e.g. ATAC-seq), and therefore do not provide a generally applicable framework for covariation-based analysis.

Here, we present Coordinate Covariation Analysis (COCOA), a method for annotating epigenetic variation across individuals using region sets. COCOA offers several advantages compared to existing methods: First, COCOA provides a flexible framework that supports both supervised and unsupervised analysis. Second, for supervised analysis, COCOA leverages covariation information by allowing continuous sample phenotypes as well as discrete groups. Third, COCOA incorporates epigenetic signal values instead of using binarized values (i.e. significant or not significant), further taking advantage of the covariation information. Finally, COCOA works with any epigenetic data that have a numerical value associated with genomic coordinates, such as DNA methylation data, chromatin accessibility data, or even multi-omics data. Importantly, no such tool that leverages covariation of epigenetic signal across samples to annotate epigenetic variation previously existed for DNA methylation data. To demonstrate COCOA’s utility, we applied it in three unsupervised analyses with DNA methylation, ATAC-seq, and multi-omics data, and a supervised analysis of DNA methylation and cancer stage. We found that across multiple data types and biological systems, COCOA is able to identify promising biological sources of epigenetic heterogeneity across sample populations.

## Results and Discussion

### An overview of COCOA

COCOA is an approach to understanding epigenetic variation among samples. COCOA derives its annotation power from a database of region sets that are grouped by function. This choice is rooted in the observation that a single effector, such as a transcription factor, often regulates many regions across the genome. Because the regions are coregulated, their epigenetic signal may covary across samples according to the activity of the effector (Fig. 1A), which can then be used to infer activity of the effector (Fig. 1B). This principle of covariation of coregulated loci or genes has been leveraged by other methods related to gene regulation^[2,3,9–13]^. To distinguish small differences among samples in the activity level of the effector, COCOA boosts statistical power by aggregating signal in region sets^[3]^.

**Figure 1.**
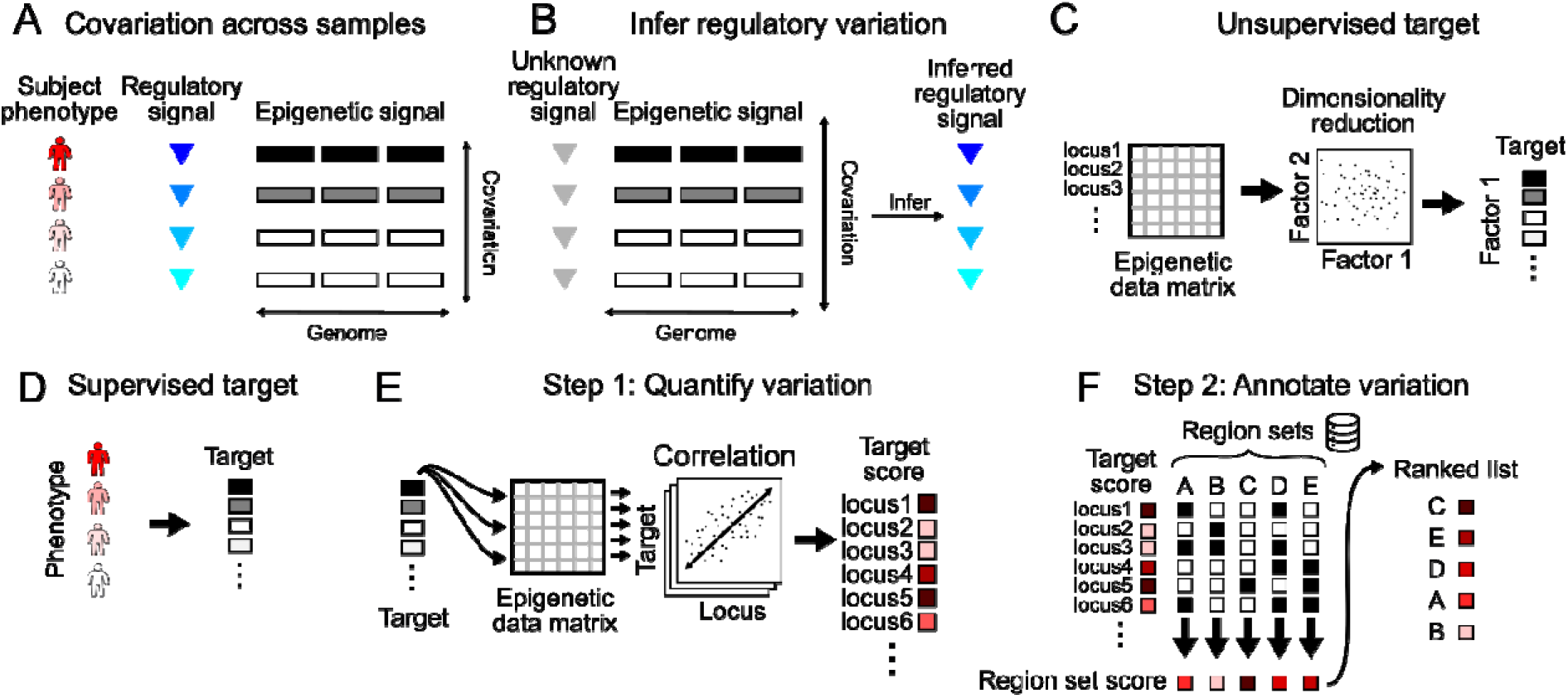
Overview of COCOA. A. A regulatory signal may covary with the epigenetic signal in the genomic regions it regulates. B. Covariation of the epigenetic signal in coregulated regions across individuals can be used to infer variation in the regulatory signal. C. COCOA can be used with an unsupervised target variable (latent factor), or D. with a supervised target variable (phenotype). E. The first step is to quantify the relationship between the target variable and the epigenetic data at each locus, resulting in a score for each locus. F. The second step is to annotate variation using a database of region sets. Each region set is scored to identify the region sets most associated with covariation between the epigenetic signal and the target variable. These top region sets can yield insight into the biological significance of the epigenetic variation.

COCOA uses this aggregated region set approach to annotate the underlying source of epigenetic variation that relates to a “target variable,” which can be either a supervised variable, like the phenotype of interest (Fig. 1C), or an unsupervised variable, like the primary latent factors in the data (Fig. 1D). COCOA annotates the inter-sample variation in the target variable by identifying region sets with variation patterns in epigenetic data that match the variation in the target variable. After a target variable is chosen, COCOA analysis consists of two main steps: first, for each locus, it computes the association of the inter-sample epigenetic variation with the target variable (Fig. 1E) and, second, it uses those associations to score a database of region sets (Fig. 1F). COCOA uses a permutation test to evaluate the statistical significance of each region set score. The result is a list of region sets ranked by how well the epigenetic signals in the region set correlate with the target variable. Highly scoring region sets have epigenetic signal that covaries across patients in the same way as the target variable, tying the functional annotation of the region set to the observed phenotypic variation.

### COCOA annotates inter-sample variation in breast cancer DNA methylation data

We first evaluated COCOA in an unsupervised analysis to determine if COCOA could identify and annotate a driving source of variation. We applied COCOA to DNA methylation data from breast cancer patients in The Cancer Genome Atlas (TCGA). In breast cancer, estrogen receptor (ER) status is a major prognostic factor and is known to be associated with a specific DNA methylation profile^[14,15]^. We first used Principle Component Analysis to identify the top four Principal Components (PCs), which we used as the target variables, and asked whether COCOA would be able to identify ER as an important source of inter-sample variation using only the DNA methylation data, without requiring the samples’ ER status.

COCOA identified a strong ER-associated signature for Principal Component 1 (PC1). This signature included many ER-binding region sets as top hits, indicating that variation of the DNA methylation in these ER-binding regions is associated with PC1 (Fig. 2A, Additional file 1: Table S1). We also identified variation in region sets for FOXA1 and GATA3, which are known to be associated with ER status^[14,15]^ (Additional file 1: Table S1). Furthermore, COCOA found the ER-associated histone modification H3R17me2 among the top scoring region sets^[16]^ (Additional file 1: Table S1). When we test the association of each PC with ER status, PC1 scores have a highly significant association with ER status (p < 10^-46^, Wilcoxon rank-sum test), whereas PC2 and PC3 are less associated (Fig. 2B). Therefore, COCOA clearly identified ER-related variation as relevant for the primary axis of inter-sample variation, despite not having access to ER status information. We found that PC4 was also associated with ER status to a lesser extent (p < 10^-20^). For PC4, COCOA identified regions with repressive chromatin marks, including binding sites for polycomb components EZH2 and SUZ12 and repressive histone modifications H3K27me3 and H3K9me3 (Fig. 2A, Fig. S1). Previous studies have linked polycomb expression to breast cancer: higher EZH2 expression is associated with ER- breast cancer^[17,18]^, EZH2 interacts with the repressor of estrogen activity (REA) protein^[19]^, and Suz12 binding sites have DNA methylation differences between ER+ and ER- breast cancer^[14]^. Therefore, PC4 represents an additional aspect of ER- related epigenetic variation. PC2 and PC3 had weaker associations with ER status (p < 0.01 and p < 10^-4^ respectively); for PC3, the highest-ranking PC3 region sets include some ER-related region sets along with hematopoietic region sets (Additional file 1: Table S1). The hematopoietic region sets may represent inter-sample variation in the immune component of the tumors since breast cancer subtypes have been reported to be associated with differing immune cell profiles^[20]^. In summary, these results demonstrate that COCOA was able to identify relevant sources of inter-sample variation without requiring known sample groups and therefore reveal COCOA’s usefulness for unsupervised analysis of DNA methylation data.

**Figure 2.**
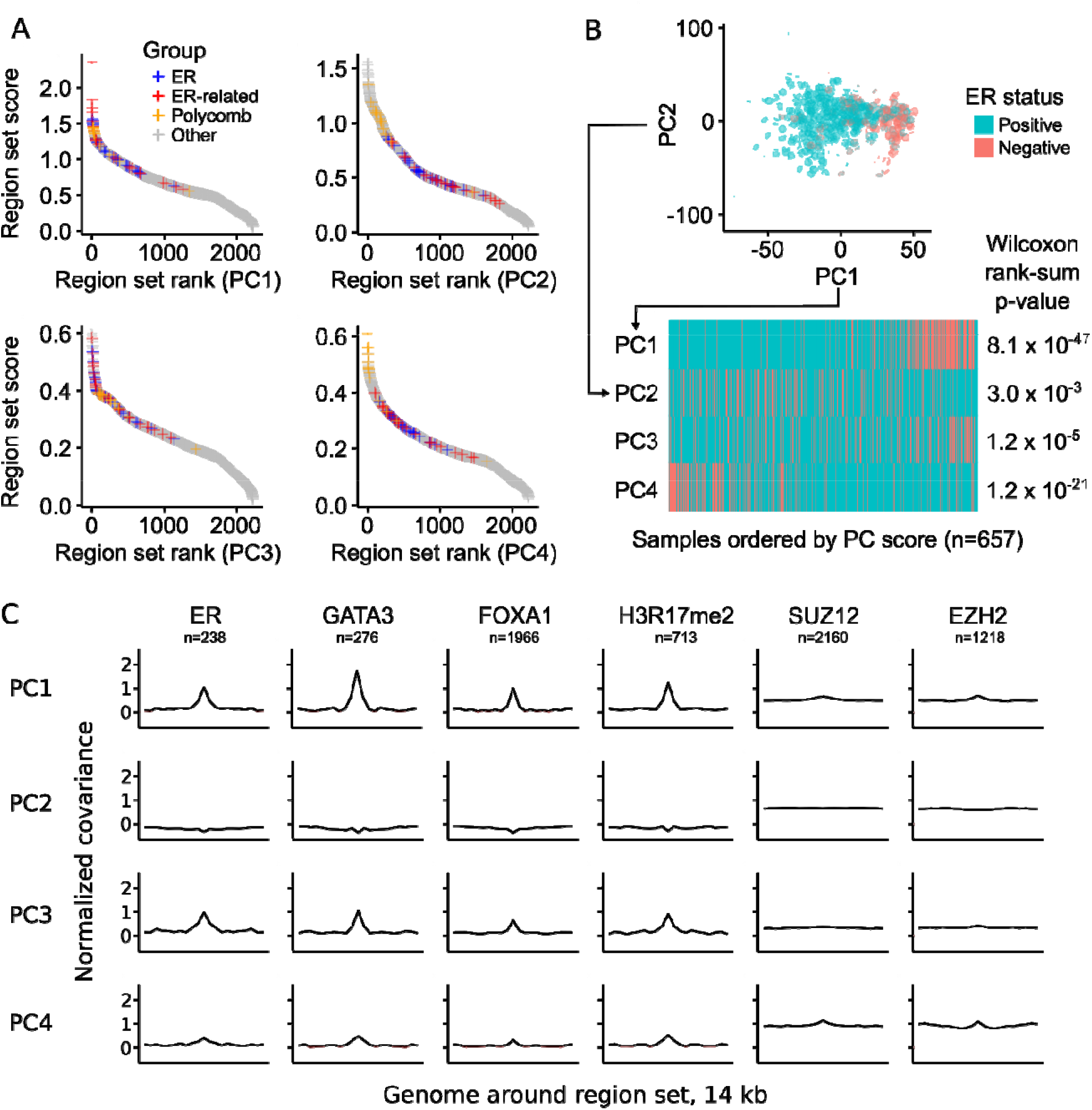
COCOA identifies sources of DNA methylation regulatory variation. A. The COCOA score for each region set, ordered from highest to lowest. The ER-related group includes GATA3, FOXA1, and H3R17me2. The polycomb group includes EZH2 and SUZ12. B. The association of PC scores with ER status for PCs 1-4 based on a Wilcoxon rank-sum test. C. Meta-region profiles of several of the highest scoring region sets from PC1 (GATA3, ER, H3R17me2) and two polycomb group proteins (EZH2, SUZ12). Meta-region profiles show covariance between PC scores and the epigenetic signal in regions of the region set, centered on the regions of interest. A peak in the center indicates that DNA methylation in those regions covaries with the PC specifically around the sites of interest. The number of regions from each region set that were covered by the epigenetic data in the COCOA analysis (panel A) is indicated by “n”.

To visualize the inter-sample variation that drives the top region sets identified by COCOA, COCOA can also plot DNA methylation in a region set, ordered by PC value (Fig. S2). Using this approach, we visualized how the DNA methylation in ER-related regions varies along PC1, demonstrating clear covariation across regions that drives the region set rankings (Fig. S2). To further confirm the specificity of the region sets, COCOA can also plot variation in broader genomic regions around the regions of interest. We found that the DNA methylation close to the transcription factor binding regions shows stronger covariation with the PC score than DNA methylation in the surrounding genome (Fig. 2C). This visualization of specificity of the covariation to the binding regions provides additional evidence of association between the PC and region set. Other high-ranking transcription factors also showed this specificity (Fig. 2C, Fig. S3). Some histone modifications, such as H3K9me3 and H3K27me3, where DNA methylation levels had high covariation with the PC showed broader regions of elevated covariation (Fig. S3). Overall, these visualization functions reveal aspects of epigenetic variation in the top region sets that could not be captured by a single region set score.

### COCOA annotates regulatory variation in ATAC-seq data

Next, we asked whether COCOA could be applied to ATAC-seq data. Unlike DNA methylation data, which annotates individual nucleotides, ATAC-seq data is summarized by accessibility values at “peak” regions ^[21]^. COCOA handles either data type. To demonstrate the region-type analysis, we ran COCOA with ATAC-seq data from TCGA breast cancer patients^[21]^, expecting that ER-related region sets would be among our top results, similar to the DNA methylation data. As before, we used PCA on the ATAC-seq data and then applied COCOA to annotate the sources of variation for each PC. We identified many of the same region sets to be associated with epigenetic variation, despite far fewer samples (657 vs 73). We found ER-related region sets to be among the top ranked results for PC1 (Fig. 3A, Additional file 1: Table S2). PC2 was characterized by high-ranking hematopoietic transcription factors (Fig. 3A, Additional file 1: Table S2), once again potentially representing inter-sample variation in the immune component of the tumors^[20]^, as in PC3 of the DNA methylation data. A few other top PCs including PC4 also had high-ranking hematopoietic transcription factors (Fig. 3A, Fig. S4). Consistent with our results, visual inspection of the chromatin accessibility signal in top ER-related and hematopoietic region sets also revealed correlation between the signal and PC scores for the PCs in which the region sets were highly ranked (Fig. S5). Polycomb region sets did not rank as prominently for the ATAC-seq data as for the DNA methylation data but there were several polycomb region sets in the top 10% of region set scores for PC4 (Fig. S6, Additional file 1: Table S2). These results are consistent with variation in ER status, which is significantly associated with PC1 and PC2 (p < 0.01, Wilcoxon rank-sum test, Fig. 3B) and to a lesser extent PC4 (p < 0.05). Visualization of the correlation between each PC and the ATAC-seq signal in the top region sets also shows specificity to the transcription factor-binding regions compared to the surrounding genome (Fig. 3C). Thus, COCOA can identify meaningful sources of variation in ATAC-seq data, providing a novel tool for regulatory analysis of ATAC-seq data.

**Figure 3.**
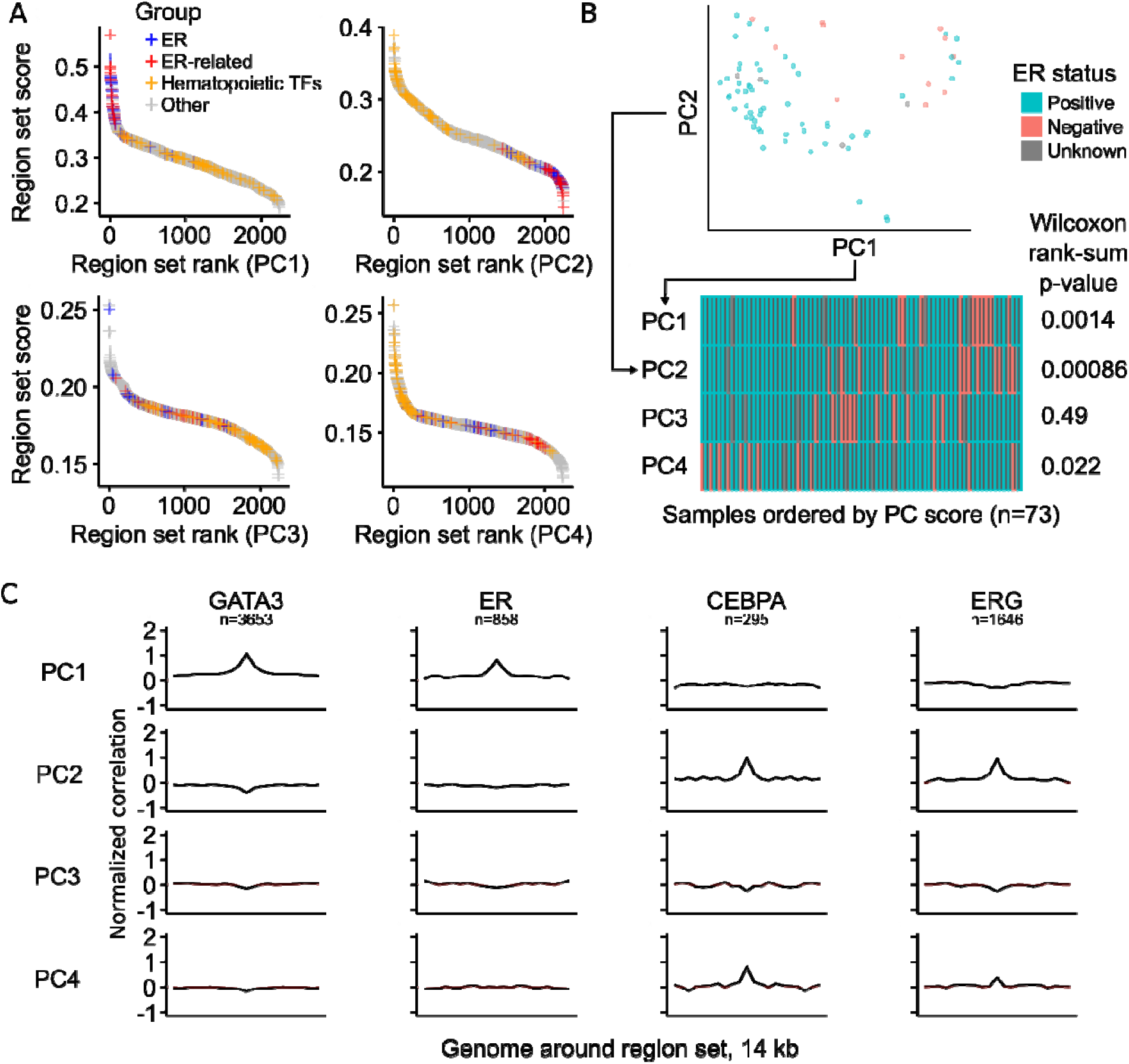
COCOA can be used for region-based data such as ATAC-seq. A. The COCOA score for each region set, ordered from highest to lowest. The ER-related group includes GATA3, FOXA1, and H3R17me2. For definition of the hematopoietic TF group, see “Region set database” in methods. B. The association of PC scores with ER status for PCs 1-4 based on a Wilcoxon rank-sum test. C. Meta-region profiles of the two highest scoring region sets from PC1 (GATA3, ER) and PC2 (CEBPA, ERG). Meta-region profiles show correlation between PC scores and the epigenetic signal in regions of the region set, centered on the regions of interest. A peak in the center indicates that chromatin accessibility in those regions correlates with the PC specifically around the sites of interest. The number of regions from each region set that were covered by the epigenetic data in the COCOA analysis (panel A) is indicated by “n”.

### COCOA identifies regulatory variation in multi-omics integration

We also aimed to determine if COCOA could annotate inter-sample variation in multi-omics analyses that integrate epigenetic data with other data types. We therefore applied COCOA to a cohort of 200 chronic lymphocytic leukemia patients^[22]^ with gene expression, *ex vivo* drug response, somatic mutation, and DNA methylation data. We used preprocessed data from MOFA (Multi-Omics Factor Analysis), a multi-omics dimensionality reduction method that summarized the high-dimensional data into 10 new dimensions referred to as latent factors (LFs)^[23]^. As part of the published analysis interpreting the 10 latent factors, the authors used a gene-centric method to annotate the latent factors with gene sets but only 5 could be associated with gene sets^[10,23]^. Because COCOA works with data associated with genomic coordinates, we were able to use the DNA methylation data from the MOFA analysis with COCOA to annotate the latent factors with region sets. Since only a subset of the DNA methylation data was used for the MOFA calculations, we calculated the correlation of each CpG in the 450k microarray with each latent factor and used this matrix as input for COCOA. Using COCOA, we are able to annotate 4 of the 5 latent factors that were not associated with gene sets, demonstrating that COCOA’s region-centric approach complements the gene-centric approach applied by the MOFA authors (Fig. 4A). For latent factor 1 (LF1), we found variability in region sets for hematopoietic regulatory regions and transcription factors (Additional file 1: Table S3), consistent with the conclusions of the original paper that LF1 is related to the hematopoietic differentiation state of the leukemic cell of origin. The top region set for LF1 was enhancer regions in the GM12878 transformed B-lymphocyte cell line, which had stark differences in DNA methylation across samples that correlated with IGHV mutation status, a marker of mature B cells that have undergone somatic hypermutation^[24]^ (Fig. 4B). This result shows that COCOA was able to identify a plausible source underlying the latent factor, a result which was not identified using gene sets. As another example, we found region sets related to stem cell biology, including OCT4, NANOG, H3K4me1 from the H9 stem cell line, and SOX2, to be associated with LF8 (Fig. 4C, Additional file 1: Table S3). Since OCT4 and NANOG activity has been shown to be associated with β catenin^[25–27]^, a mediator of WNT signaling, our results support and further expand upon the original association between LF8 and WNT reported by the MOFA authors. These results demonstrate that COCOA can enable richer multi-omics analysis by annotating the epigenetic component of inter-sample variation.

**Figure 4.**
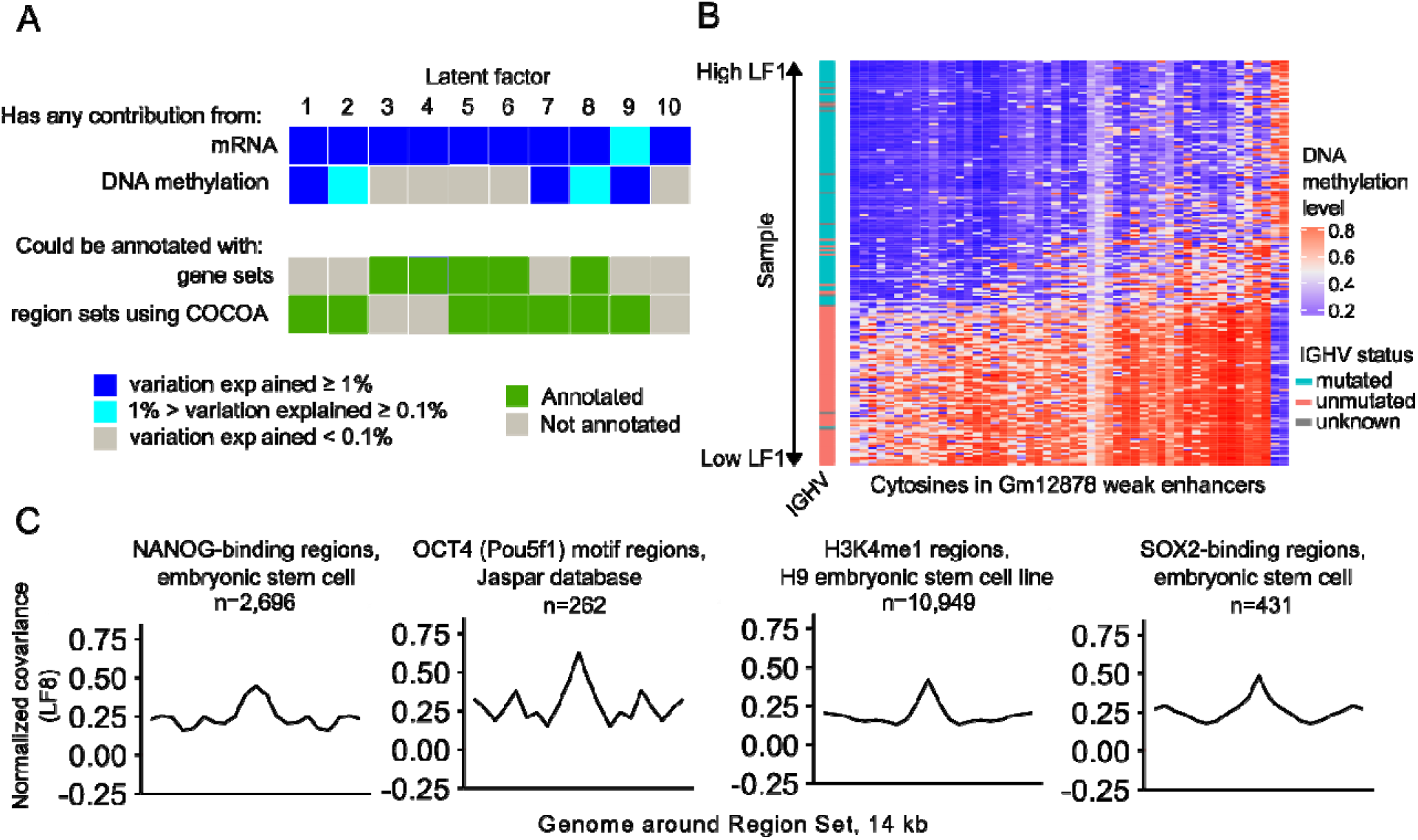
COCOA can be applied to multi-omics analyses that include epigenetic data. A. COCOA can annotate latent factors that were not annotated by a gene set approach. In the top of panel A, dark blue indicates that the data type explained at least 1% of the variation of the latent factor while light blue indicates that the data type explained between 0.1% and 1% of the variation. Gray indicates less than 0.1% explained. In the bottom of panel A, green indicates that at least one statistically significant gene set or region set was found for the latent factor and gray indicates no significant gene or region sets were found. B. COCOA identifies an enhancer region set from a transformed B-lymphocyte cell line where DNA methylation is correlated with latent factor 1 and IGHV mutation status, a marker of mature B cells that have undergone somatic hypermutation. The 50 CpGs with the highest absolute correlation with LF1 from the region set are shown. C. Meta-region profiles show covariation between DNA methylation and LF8 score in certain regions bound by transcription factors functional in stem cell biology and by H3K4me1 in a stem cell line compared to the surrounding genome. The number of regions from each region set that were covered by epigenetic data in the COCOA analysis is indicated by “n”.

### COCOA reveals associations between epigenetic state and variation in sample phenotype

The three examples thus far demonstrate how COCOA can be applied in an unsupervised analysis, which explores biological variation in the absence of known groups. To explore whether we could apply COCOA to a setting where groups or phenotypes are known, we extended COCOA to accommodate supervised analysis. For the supervised approach, we select a sample phenotype of interest (such as a molecular phenotype or a clinical outcome) and then measure the association of epigenetic variation with that parameter. To demonstrate a supervised COCOA analysis, we analyzed TCGA 450k methylation microarrays from kidney renal clear cell carcinoma (KIRC). This dataset includes a phenotypic annotation of cancer stage, which we used as our target variable. We hypothesized that COCOA could associate an epigenetic regulatory state with cancer stage and decreased survival. To test this hypothesis, we used COCOA to identify region sets where DNA methylation is correlated with cancer stage. We used a training-validation approach to assess significance of our results (see Methods). In the training samples, COCOA identified polycomb protein (EZH2 and Suz12)-binding region sets to have the highest correlation with cancer stage (Fig. 5A, Additional file 1: Table S4). Next, we tested whether the average DNA methylation level in the top EZH2 region set is associated with cancer stage. In both training and validation samples, average DNA methylation level in EZH2-binding regions had a significant positive correlation with cancer stage (p < 10^-6^ and p < 10^-7^, *t* approximation) showing that the COCOA result extends beyond the training set (Fig. 5B, Additional file 1: Table S5). Higher DNA methylation levels in EZH2-binding regions in advanced stages of cancer suggest that these regions could be repressed in advanced cancer stages, which would be consistent with higher activity of the repressive protein EZH2. This result is consistent with previous studies, which have found that higher EZH2 expression could promote metastasis in renal cell carcinoma^[28]^ and other cancers^[29,31]^ and is associated with a more advanced cancer stage^[32,33]^.

**Figure 5.**
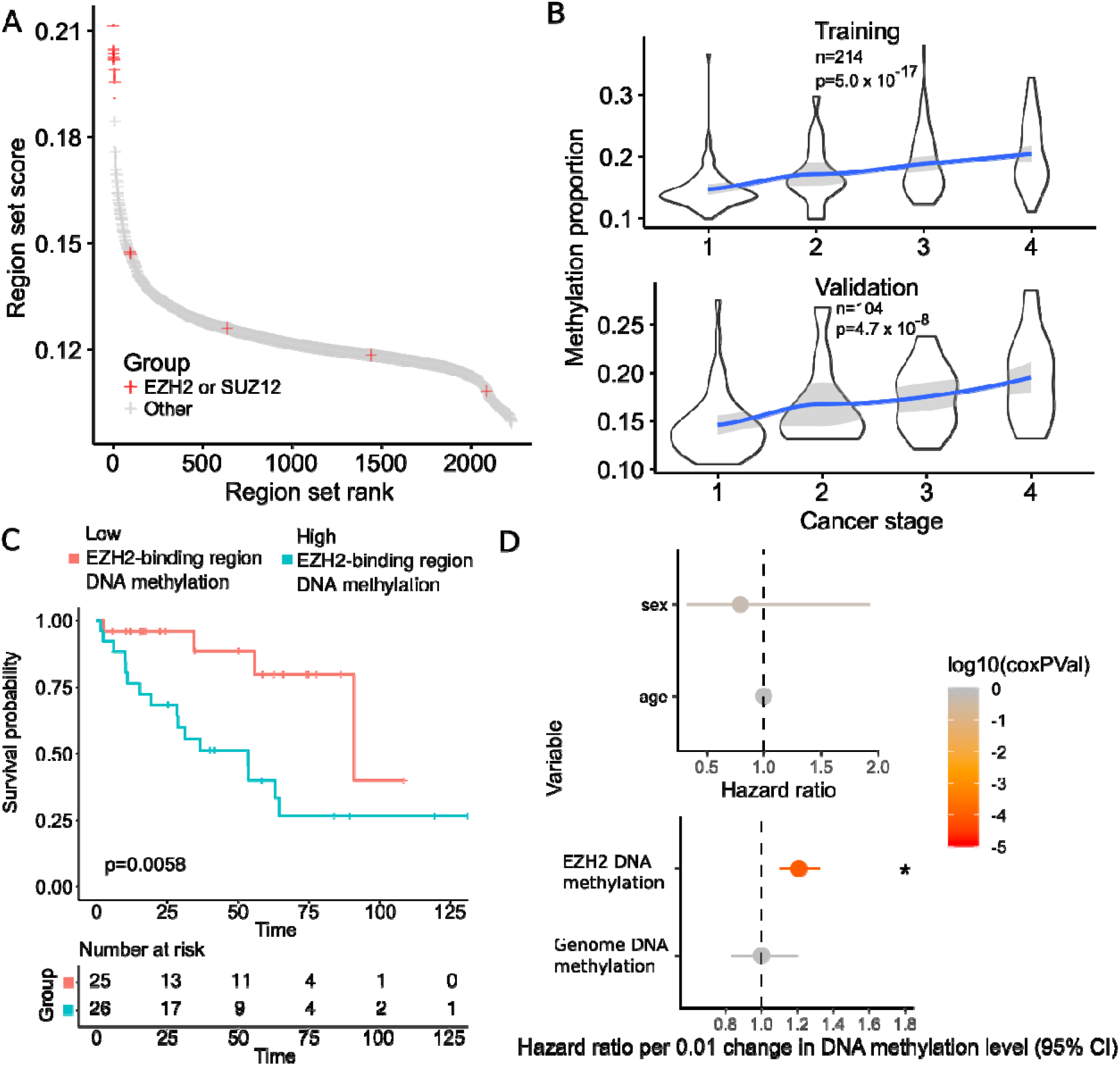
COCOA identifies region sets related to a patient phenotype of interest, cancer stage. A. Region sets for the polycomb proteins EZH2 and SUZ12 were the top region sets related to cancer stage. B. Average DNA methylation level in EZH2-binding regions (the top EZH2 region set) increases with cancer stage. P-values by t approximation with null hypothesis that correlation is zero. C. Kaplan-Meier curves of the validation samples, grouping samples by average DNA methylation in EZH2 binding regions (25% highest samples and the 25% lowest samples). P-value from log-rank test. E. Cox proportional hazards model of average DNA methylation in the top EZH2-binding region set, correcting for age, gender, and average genome methylation level.

To further assess the relevance of our COCOA results, we tested the association between DNA methylation in our top EZH2 region set and patient survival. We compared the quartile of patients with highest average DNA methylation in the EZH2 region set to the quartile of patients with the lowest average DNA methylation, using a Kaplan-Meier estimate (Fig. 5C). Patients with higher EZH2 region set DNA methylation have significantly decreased survival compared to those with lower DNA methylation (p < 0.01, log-rank test, Fig. 5C). A Cox proportional hazards model correcting for age, gender and average genome methylation levels also revealed a significant association between average DNA methylation level in EZH2 binding regions and patient survival (p < 10^-4^, Fig. 5D, Additional file 1: Table S6). Previous studies found EZH2 expression to be prognostic for survival in renal cell carcinoma and other cancers^[29,32,34]^, but to our knowledge, this is the first demonstration that DNA methylation levels in EZH2 binding regions could be prognostic for survival in renal cell carcinoma. We further assessed for cancer stage and survival association for the top TF region sets from the COCOA analysis. We tested the two highest scoring TFs - JUND and TCF7L2. In the validation data, DNA methylation in JUND-binding regions had a significant negative correlation with cancer stage (p=0.022, *t* approximation) but we could not validate its association with survival because it did not satisfy the Cox proportional hazards assumption (Fig. S7A, Additional file 1: Table S6). DNA methylation in TCF7L2-binding regions was not significantly correlated with cancer stage in the validation data (Fig. S7B, Additional file 1: Table S5) but higher DNA methylation was significantly associated with better overall survival (p= 0.038, Cox proportional hazards model, Fig. S7C, Additional file 1: Table S6). Through this supervised analysis, we demonstrate that COCOA can identify epigenetic variation related to a given sample phenotype of interest, providing a novel means for targeted analysis of epigenetic variation.

### DNA methylation in EZH2-binding regions is associated with cancer stage and survival in multiple cancers

Given that COCOA identified associations with EZH2 region sets in both our unsupervised analysis of breast cancer and our supervised analysis of kidney renal cell carcinoma, we wondered whether the link with EZH2 and DNA methylation would hold true for other cancer types. To test this, we performed a pan-TCGA analysis investigating the association between average DNA methylation in EZH2/SUZ12- binding regions and cancer stage as well as overall patient survival. We combined regions from the top group of 11 EZH2 and SUZ12 region sets from the KIRC analysis (Fig. 5A, Additional file 1: Table S4) to generate a single EZH2/SUZ12 region set, referred to hereafter simply as EZH2-binding regions. We then computed the average DNA methylation in this region set for each sample and tested its association with either cancer stage or overall survival. We found a significant correlation between DNA methylation in EZH2-binding regions and cancer stage in multiple cancer types (Fig. S8, Additional file 1: Tables S7 and S8). DNA methylation in EZH2-binding regions positively correlated with cancer stage in 5 of 21 tested cancers, but trended negative in 3 cancer types (Fig. S8), of which colon adenocarcinoma (COAD) had a significant negative correlation (p < 0.05, *t* approximation, Holm-Bonferroni correction), consistent with a previous report^[35]^. To further investigate the significance of the EZH2-binding regions, we used a Cox proportional hazards model to test for association between survival and average DNA methylation in these regions and found a significant association in 5 cancer types (Fig. 6, Additional file 1: Tables S8 and S9). Similar to the cancer stage analysis, higher DNA methylation level was more often associated with increased risk of death, but trended to lower risk in a few cancer types (Fig. 6). This result is consistent with previous reports that EZH2 can be either oncogenic or a tumor suppressor^[31,36,37]^ and emphasizes the context-specific effects of EZH2. Our pan-cancer analysis also supports previous reports suggesting that polycomb activity may be commonly dysregulated in cancer^[31]^ and may influence survival in a variety of cancers, with some cancers having a positive and others a negative association^[31]^. Our results contrast with previous reports for several cancer types (Supplementary Discussion). This analysis identified a novel connection between EZH2 and survival in adrenocortical carcinoma (ACC), which has not been previously demonstrated. Furthermore, we have shown for the first time that variation in DNA methylation at EZH2-binding regions is associated with cancer stage and patient survival across a variety of cancers. Overall, this analysis demonstrates the ability of COCOA to annotate epigenetic variation and its potential to generate new mechanistic hypotheses about epigenetic heterogeneity and disease drivers.

**Figure 6.**
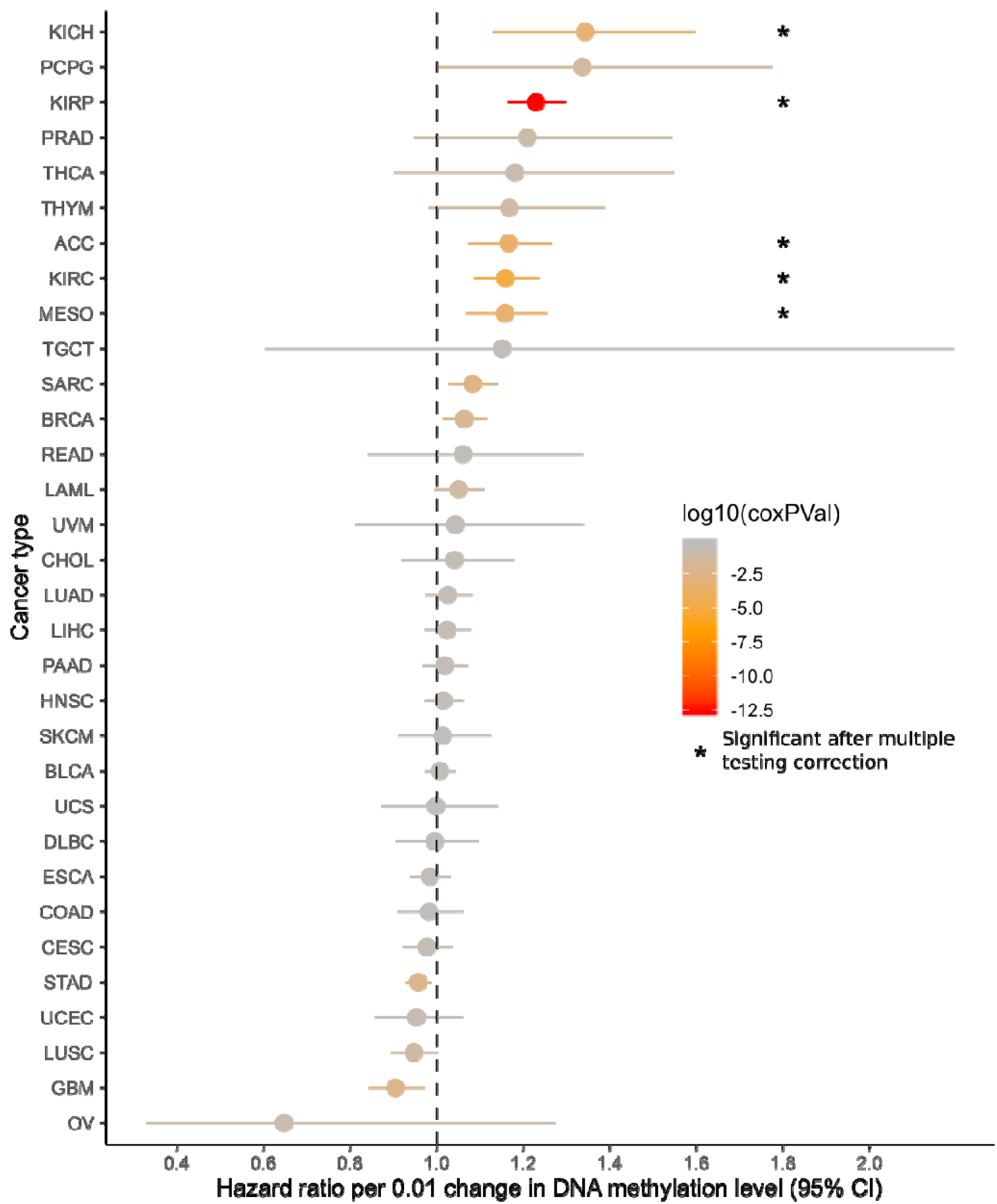
Pan-cancer survival analysis of DNA methylation in EZH2/SUZ12-binding regions. The mean hazard ratio and 95% confidence interval for the average DNA methylation in EZH2/SUZ12-binding regions are shown for each cancer type. Color indicates the raw p-values and asterisks mark significance after Holm-Bonferroni correction.

### Comparison of COCOA to other methods

COCOA distinguishes itself from other methods by being the only method of its type for DNA methylation data and by its flexibility in supporting a wide range of analyses for epigenetic data. We conceptualize COCOA as being in a class of methods that relies on covariation of epigenetic signal to annotate epigenetic variation. This separates COCOA from the methods that annotate epigenetic variation without taking into account covariation. To demonstrate the power of this approach, we compared COCOA to LOLA, a method that does not consider covariation. This analysis demonstrated that COCOA has superior ability to mitigate noise (Supplemental Information; Fig. S9; Additional file 1: Tables S10, S11, S12). Other methods that do take into account covariation have key differences from COCOA. First, while tools exist that aggregate signal in related groups such as gene sets or region sets and use PCA to identify covariation of signal across samples (Table 1), no existing tool does this for DNA methylation data. Second, COCOA creates a generalized framework for region set analysis which results in great flexibility in applications. This generalized framework allows COCOA to be used in analyses that other tools may not support: with multiple epigenetic data types, for supervised or unsupervised analyses, with a variety of mathematical metrics, and for single-omic or multi-omic analyses. For a brief description of each method from Table 1 and further comparison to COCOA, see “Comparison of COCOA to other region set or covariation-based methods” in the supplementary text. Of the epigenetic tools with similar goals to COCOA, chromVAR^[2]^ is the most widely used and most similar to COCOA in its input type. Therefore, we selected chromVAR for comparison to COCOA with the breast cancer ATAC-seq data. Each method revealed relevant but partially divergent aspects of inter-sample variation. COCOA had an improved ability to identify ER-related epigenetic variation and to separate biological signals with its use of PCA (Fig. S10, Supplementary Information: “Comparison of COCOA to chromVAR”, Additional file 1: Tables S2, S13). COCOA also extends beyond chromVAR in COCOA’s analysis options and supported data types. COCOA thus provides a novel framework for flexible covariation-based analysis of DNA methylation and other epigenetic data, ameliorate

**Table 1.**
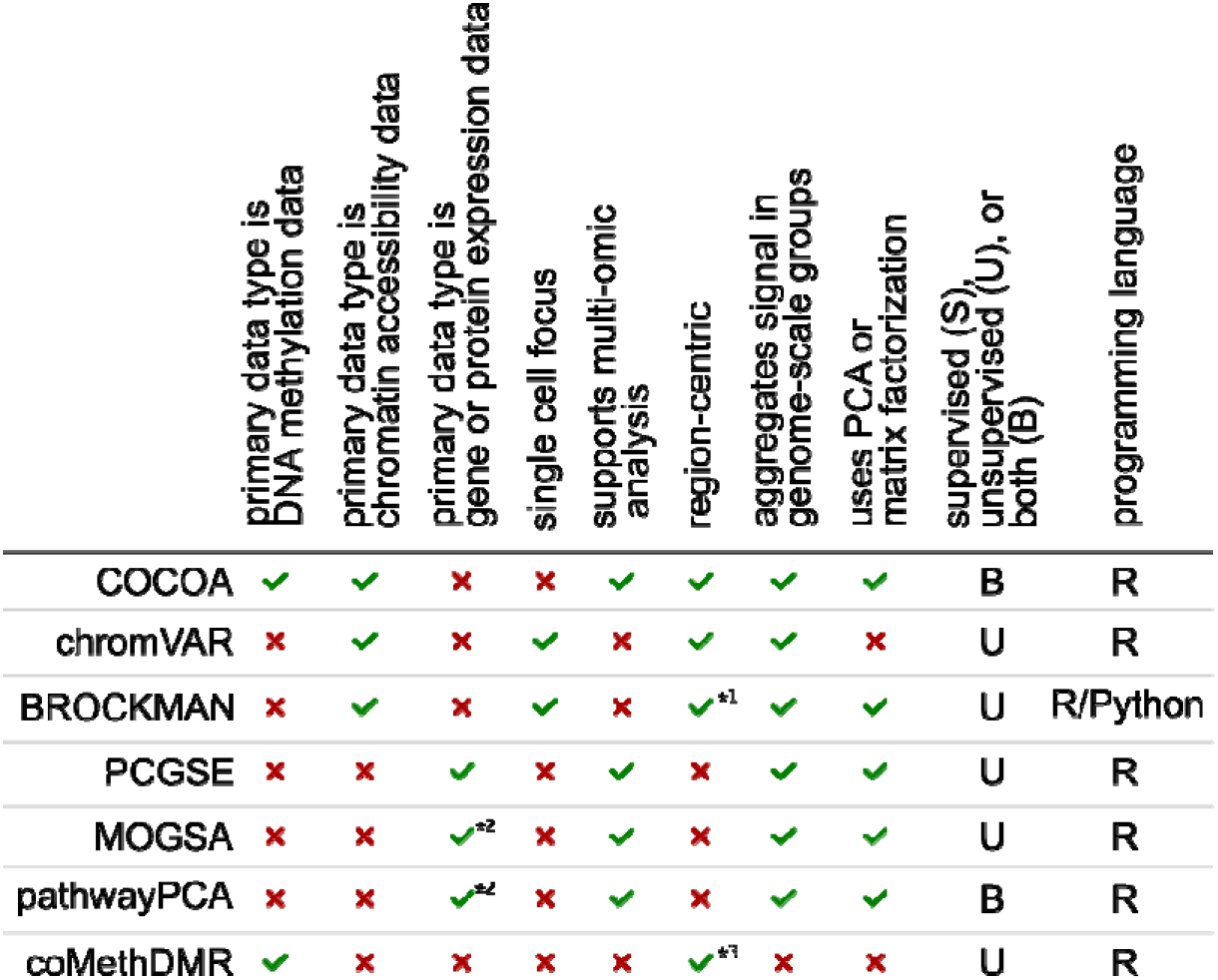
Features of COCOA and related methods. *^1^BROCKM AN uses k-mer counts but the regions containing each k-mer can be conceptualized as a region set. *^2^MOGSA and pathwayPCA can involve multiple “omics” data types but including gene-centric data such as gene or protein expression is important for the methods. *^3^coMethDMR finds differentially methylated regions but often annotates them in reference to genes.

## Conclusion

We created a flexible framework for identifying and understanding sources of regulatory variation in epigenetic data. COCOA could be applied to any epigenetic data that has a value associated with genomic coordinates, which includes both nucleotide-level data such as bisulfite sequencing and region-based data such as ATAC-seq data. Our results also demonstrate how COCOA can be integrated with multi-omics analyses that include epigenetic data. Our tool allows scientists to leverage publicly available regulatory data to annotate variation in their epigenetic data. In an unsupervised analysis, COCOA can annotate the major axes of inter-sample variation. In a supervised analysis, COCOA can annotate inter-sample variation related to a specific phenotype of interest. We have released COCOA as a Bioconductor package^[38]^, facilitating this new method of regulatory analysis. COCOA is a flexible and powerful method for interpreting regulatory variation between individuals.

## Methods

### COCOA algorithm

#### Overview

COCOA annotates variation in epigenetic data through two steps. In the first, we quantify the association between each feature in the epigenetic data and the target variable using a metric such as correlation (Fig. 1C). This gives a score to each epigenetic feature that represents how much it is associated with the target variable. Then in the second step, we use the epigenetic feature scores to score region sets from a large collection of region sets (Fig. 1D). Finally, we use a permutation test to assess statistical significance, and return a ranked list of region sets.

#### Step 1: Quantifying variation across samples

COCOA starts with a data matrix of epigenetic signal values in genomic regions, where each row is a genomic locus (e.g. a CpG or an ATAC-seq region), and each column is a sample. The values in the matrix correspond to signal intensity levels (e.g. DNA methylation level or chromatin accessibility) of a given sample at a given locus. The first step in a COCOA analysis is to transform the original data into a score for each locus measuring how much it contributes to the target inter-sample variation. We refer to the score for an epigenetic feature (locus) as a “feature contribution score” (FCS). This calculation can be either supervised or unsupervised (Fig. 1B):

##### Supervised

For supervised analyses, the goal is to identify sources of variation associated with a target sample phenotype of interest. Therefore, in addition to the epigenetic data matrix, we require a vector representing the target sample phenotype. We then quantify the association between the target sample phenotype and the epigenetic signal at each genomic locus using a method such as Pearson correlation. We end up with a vector of scores (which for correlation is the correlation coefficient) representing how strongly epigenetic variation at a genomic locus is associated with variation in the sample phenotype. Metrics other than Pearson correlation can be used to quantify variation, as long as they produce a score for each genomic locus. A detailed discussion of metric choice follows in the section, *Metric for quantifying variation.*

##### Unsupervised

For unsupervised analyses, we first apply a dimensionality reduction technique such as PCA or MOFA^[23]^ to identify latent factors that represent significant sources of inter-sample variation^[10]^. Then, we treat these latent factors as target sample phenotypes and quantify the association between each latent factor and the epigenetic data as we would for the sample phenotype in the supervised analysis. In this case, the feature contribution score for each genomic locus represents how strongly epigenetic variation at that genomic locus is associated with variation in the latent factor.

#### Step 2: Annotate variation with the COCOA algorithm

After quantifying inter-sample variation, we are left with one or more vectors that assign FCS to each genomic locus in the original data matrix. COCOA next seeks to determine which region sets are associated with that variation. For this step, COCOA relies on a database of region sets. Here, we have used a subset of the LOLA database^[1]^, which includes several thousand region sets that have been manually collected from several large-scale experiments and databases, including the ENCODE^[39,40]^ and Roadmap Epigenomics projects^[41,42]^. For the sample-specific data, COCOA can operate on two types of signal data: single-nucleotide data (e.g. DNA methylation) or region-based data (e.g. ATAC-seq peaks). In either case, we will aggregate the scores for all individual genomic loci into a combined score for each region set (Fig. 1D). Due to different experiments testing the same TF or histone modification, some region sets share similar regions to each other and therefore their scores are not completely independent.

For single base-pair resolution data (e.g. DNA methylation data), the following algorithm is used for a single region set and a single FCS vector: First, we optionally take the absolute value of the FCS (Supplementary Methods). Then, we identify all features whose genomic coordinates overlap the given region set. Within each region from the region set, we average the FCS of any overlapping features to get a single average value for each region. We then average the region scores to get the final score for that combination of region set and FCS vector. This score represents how much that region set is associated with the latent factor or phenotype that corresponds to the FCS vector. We repeat this process for each pairwise combination of region set and latent factor/phenotype FCS vector.

For region-based data such as ATAC-seq data, the scoring is conceptually similar to single-nucleotide data, but with slight differences. We use the following algorithm: To score a region set for a given latent factor or phenotype, we first identify all overlaps between “data regions” (regions for the epigenetic signal data) and region set regions. For each overlap, we calculate what proportion of the region set region is overlapped by the data region. We then take a weighted average of the FCS of all the overlapping data regions, weighting each data region’s FCS by the proportion that region overlaps a region set region and dividing by the sum of all overlap proportions. This weighted average is the region set score that represents how much the region set is associated with the latent factor or phenotype. We repeat this process for each combination of region set and latent factor/phenotype.

COCOA also offers alternative scoring methods including the option to use the median instead of the mean. We discuss this option in the Supplementary Discussion where we compare results for COCOA of breast cancer DNA methylation using median and mean scoring methods, finding overall similar results and high correlation between median and mean scores (Fig. S11, Additional file 1: Tables S1, S14). Other scoring options can be found in the software documentation.

#### Metric for quantifying variation

Choosing an appropriate metric can help to effectively capture the relationship between epigenetic variation and variation in the target variable (Supplementary Methods). In this paper, we used covariance, Pearson correlation, Spearman correlation, PCA, and MOFA^[23]^ to quantify variation, but other variation metrics and dimensionality reduction techniques can be used with COCOA for quantifying inter-sample variation, depending on the specific circumstances of a given analysis. The only requirement is that the metric must provide a score for each epigenetic locus that quantifies how much it is associated with variation in the target variable. The choice of metric can depend on the data type.

For DNA methylation data, since DNA methylation data is bounded from 0 to 1, we used covariance to give greater weight to CpGs with larger changes in DNA methylation across samples. Since the range of ATAC-seq counts could be very different between different peaks, we used Pearson correlation for the ATAC-seq data in order to give each peak a comparable score, regardless of the peak’s range. This principle also applies to PCA. When performing PCA, we recommend scaling the data by dividing each variable by its variance for ATAC-seq data (equivalent to correlation) but not for DNA methylation data (equivalent to covariation). Then, when treating the principal components as the target variables, we use the corresponding metric -- covariance or correlation -- to get the feature scores. We recommend Spearman correlation when the relationships between the target variable and the epigenetic features are monotonic but not linear, as may occur when the target variable is ordinal (e.g. cancer stage).

### Permutation test

To assess statistical significance of the COCOA results, we use a permutation test. For both supervised and unsupervised COCOA analyses, we have a target variable (i.e. the sample phenotype or latent factor) and want to understand the relationship between the target variable and the epigenetic data. For a single permutation, we randomly shuffle the samples’ target variable values then recalculate the association between the epigenetic data and the target variable as done in Step 1 (Fig. 1C). This gives each epigenetic feature an FCS for the shuffled target variable. Then we run COCOA on the new feature contribution scores to score each region set in the database. This process is repeated for each permutation. The COCOA scores for a given region set from the permutations form a region set-specific null distribution. Because the sample labels were shuffled instead of the epigenetic data, the null distributions can appropriately capture the correlation structure of the epigenetic data, accounting for the correlation between epigenetic features in a given region set. The region set-specific null distributions also protect against false positives that could arise from some region sets being more fully covered by the epigenetic assay than others because each score in a region set’s null distribution is created from the same coverage profile. To reduce the computational burden, we calculated 300 permutations and applied a permutation approximation technique^[43]^. We fit a gamma distribution to each null distribution using the method of moments in the fitdistrplus R package^[44]^ and then calculated a p-value for each region set using its gamma distribution. To test the appropriateness of fit of the gamma approximation, we ran a simulation study with 100,000 permutations, and then subsampled and applied the approximation to see how close the approximation is to the true p-value. Our conclusion is that the gamma approximation is accurate for high p-values, but the gamma approximation may overestimate the significance of low p-values; therefore, we advise that it can be helpful for screening out region sets that are not significant (Fig S12; further discussion in Supplementary Information). To correct p-values for the number of region sets tested, we used Benjamini-Hochberg false discovery rate (FDR) correction^[45]^ with an FDR of 5%.

### Meta-region profile plots

To visualize results, COCOA produces a plot we call the *meta-region profile* plot (e.g. Fig 2C). The goal of the meta-region profile is to compare the feature contribution scores in the regions of interest to the surrounding genome to assess how specific the captured signal is to a region set. We combine information from all regions of the region set into a single summary profile as has been done for DNA methylation data^[3,6]^. Each region in the region set is expanded on both sides to include the surrounding genome (e.g. expanded to 14 kb total, centered on the region of interest). This enlarged region is then split into bins of approximately equal size. Finally, the FCS for corresponding bins from each region of the region set are averaged to get a single “meta-region” FCS profile. A peak in the middle of this profile suggests that there is variation that is specific to this region set.

### Region set database

To annotate variation in the epigenetic data, we used a subset of the LOLA database^[1]^ (filtered with R script, see Supplementary Materials), totaling 2246 region sets from public sources. Sources included the ENCODE project^[39,40]^, Roadmap Epigenomics^[41,42]^, CODEX database^[46]^, and the Cistrome database^[47]^. Additionally, we included some region sets derived from JASPAR motif^[48]^ predictions. Examples of region sets include transcription factor binding sites from ChIP-seq experiments, histone modification regions from ChIP-seq experiments, and cell type or condition-specific accessible chromatin from ATAC-seq experiments. For a discussion of how to choose a region set database and other related considerations, see Additional file 1. For each analysis, we only considered in the results region sets that had at least 100 regions with any coverage by the epigenetic data. Since the CLL MOFA data was in reference genome hg19 and the breast cancer data was in hg38, we used the corresponding hg19 or hg38 version of the region set database when analyzing each dataset. A brief description of the region sets can be found in the supplementary data (Additional file 1: Tables S1-S4) and the database is available at http://databio.org/regiondb^[1]^. To designate region sets “hematopoietic TFs” for Fig. 3, we did a literature search, selecting three reviews: one focusing on myeloid TFs^[49]^, one focusing on lymphoid TFs^[50]^ and one general hematopoietic TF^[51]^. The hematopoietic TFs identified from these reviews are the following: RUNX1, TAL1, PU.1, CEBPA, IRF8, GFI1, CEBPE^[49]^, TCF3, EBF1, PAX5, FOXO1, ID2, GATA3^[50]^, KLF1, GATA1, GATA2, IKZF1, CMYB, and NFE2^[51]^. Since GATA3 was also identified as an ER-related TF, we did not consider GATA3 as a hematopoietic TF in plots to avoid confusion.

### Breast cancer analyses

#### Datasets

For the unsupervised breast cancer analyses, we used DNA methylation and ATAC-seq datasets from The Cancer Genome Atlas (TCGA). We retrieved the DNA methylation and clinical data with the TCGAbiolinks R package^[52]^. We identified 657 patients with both 450k DNA methylation data and known ER and progesterone status. For the ATAC-seq data, we retrieved a peak count matrix for the consensus set of breast cancer ATAC-seq peaks identified by Corces et al. from the following location: https://atacseq.xenahubs.net/download/brca/brca_peak_Log2Counts_dedup. We used a sample ID lookup table to match the ATAC-seq IDs to the standard TCGA identifiers: https://gdc.cancer.gov/about-data/publications/ATACseq-AWG. We excluded one patient of the 74 patients with ATAC-seq data (TCGA-AO-A0J5) for whom we did not have sufficient metadata.

#### Data processing and quantifying variation

For the breast cancer DNA methylation data, we excluded the sex chromosomes. For the ATAC-seq data, we used the peak count matrix from Corces et al.^[21]^, without further processing. We performed PCA on the DNA methylation data and the ATAC-seq data separately with the ‘prcomp’ R function, with centering and without scaling. PCA is used to get covariance of features and to prioritize the largest sources of covariance. After PCA, we calculated the covariance or correlation coefficient for each epigenetic feature with each latent factor to get a value that represented how much each feature contributed to each latent factor. We used covariation for the DNA methylation data and correlation for the chromatin accessibility data. To test the association of ER status with PC score, we used the Wilcoxon rank-sum test with ER positive samples and ER negative samples as the two groups.

#### Comparison of COCOA and chromVAR

To compare COCOA and chromVAR^[2]^, we completed two tests with the breast cancer ATAC-seq data. First, we applied chromVAR with the same region set database used by COCOA in our ATAC-seq analysis. Second, we applied COCOA and chromVAR with the main motif database used by chromVAR in its publication, which is a curated version of the cisBP database^[53]^ and is available as the “human_pwms_v1” data object from the “chromVARmotifs” R package that can be downloaded from the “GreenleafLab/chromVARmotifs” Github repository. We applied chromVAR to the normalized data from Corces et al., adding a pseudocount to bring the minimum normalized signal up to zero. To use the motif database with COCOA, we identified peaks with motif hits using the “matchMotifs” function from the “motifmatchr” R package^[54]^ with default parameters and took those regions as a region set. The “matchMotifs” function is the method chosen by chromVAR authors for identifying motif matches in the chromVAR Bioconductor vignette. For the chromVAR figure, we designated motifs as AP-1-related based on an AP-1 review (Figure 1 of review)^[55]^.

### Multi-omics chronic lymphocytic leukemia analysis

#### Datasets

For the unsupervised multi-omics analysis, we used preprocessed data that was included with the MOFA R package, specifically the latent factors from the multi-omics dimensionality reduction analysis of 200 chronic lymphocytic leukemia (CLL) patients as described by Argelaguet et al.^[22,23]^. We retrieved the 450k DNA methylation data for these patients using the ExperimentHub R package^[56]^ (CLLmethylation data package, ExperimentHub ID: EH1071)^[22]^.

#### Data processing and quantifying variation

For the multi-omics analysis, we used the dimensionality reduction results from the paper by Argelaguet et al. and then extended the results to CpGs that were not included in the dimensionality reduction. The original multi-omics analysis used only the most variable 1% of CpGs (4,248 CpGs) for calculation of the latent factors. Since COCOA benefits from higher coverage of CpGs across the genome, we calculated the correlation of each CpG from the DNA methylation microarrays (excluding sex chromosomes) with each latent factor. This yielded a matrix with CpG, latent factor correlations where each row is a CpG and each column is a latent factor, which can be used as input to COCOA.

### Kidney renal clear cell carcinoma analysis

#### Dataset

For the supervised KIRC analysis, we used DNA methylation and clinical data from The Cancer Genome Atlas. We used 450k DNA methylation microarray data for 318 patients, retrieved with the curatedTCGAData R package^[57]^. The clinical data included cancer stage and survival information that was used to label samples in the supervised analysis.

#### Data processing and quantifying variation

For the supervised analysis of KIRC methylation, we first split the data into two groups: training (2/3 of patients) and validation (1/3 of patients), keeping approximately equal proportions of each cancer stage in each group. With the COCOA samples, we first calculated the Spearman correlation between the DNA methylation levels and the sample phenotype of interest, cancer stage. This resulted in a correlation coefficient for each CpG. We then applied the COCOA algorithm on the absolute correlation coefficients.

#### Validation and survival analysis

After running COCOA on 2/3 of the samples, we did validation analyses on the remaining 1/3 of samples. First, we tested whether each patient’s average DNA methylation level in the top EZH2 region set from COCOA was correlated with cancer stage, using the ‘con.test’ R function^[58]^ and Spearman correlation. To calculate correlation p-values for the null hypothesis that the correlation was zero, we used an asymptotic *t* approximation, the default method used by the ‘con.test’ function. To calculate the average methylation, we first separately averaged DNA methylation within each EZH2 region, then averaged all the region averages. We also tested whether average DNA methylation in EZH2 regions was related to overall patient survival. We created Kaplan-Meier curves with two groups: the 25% of validation samples with highest DNA methylation in EZH2 regions and the 25% of samples with the lowest DNA methylation. We used a log-rank test from the ‘survminer’ R package’s ‘ggsurvplot’ function^[59]^ to get a p-value for the Kaplan-Meier curves. We created a Cox proportional hazards model with all validation samples, relating average DNA methylation in EZH2 regions to patient survival and correcting for age, gender, and average genome methylation level. We also tested the two highest scoring TF region sets from the COCOA analysis -JUND and TCF7L2 -- for association with cancer stage and survival using the methods described above. We tested whether variables satisfied the proportional hazards assumption using the ‘cox.zph’ function in R^[60–62]^ (Additional file 1: Table S6), considering variables with p < 0.05 as not satisfying the assumption. The JUND validation model did not meet the assumption for the variable of interest (average DNA methylation in EZH2/SUZ12-binding regions) and therefore was not considered.

#### Pan-cancer EZH2 analysis

In this analysis, we tested whether average DNA methylation level in EZH2-binding regions would be associated with cancer stage and patient survival in other cancer types than KIRC. We combined regions from the top group of 11 EZH2 and SUZ12 region sets from the KIRC analysis (Fig. 5A, Supplementary Data) to make a single “master” EZH2/SUZ12 region set (referred to as EZH2-binding regions). We took the union of all regions and merged regions that overlapped. We downloaded DNA methylation microarray data for 33 TCGA cancer types using the curatedTCGAData R package^[57]^. Then, for each sample, we calculated the average DNA methylation level in EZH2-binding regions. For each cancer type for which we had cancer stage information (21/33), we calculated the Spearman correlation between average EZH2-binding region DNA methylation and cancer stage, using the ‘con.test’ R function^[58]^. To calculate correlation p-values for the null hypothesis that the correlation was zero, we used an asymptotic *t* approximation, the default method used by the ‘con.test’ function. Next, for each cancer type, we used a Cox proportional hazards model to test the association of average EZH2-binding region DNA methylation with survival, with the covariates patient age, sex, and average microarray-wide DNA methylation level as available. We tested whether variables satisfied the proportional hazards assumption using the ‘cox.zph’ function in R^[60–62]^ (Additional file 1: Table S9). We considered variables with p < 0.01 as not satisfying the assumption, picking a more stringent cutoff because more models were tested. Models that did not meet the assumption for the variable of interest (average DNA methylation in EZH2/SUZ12-binding regions) were removed, in our case only one cancer type- low grade glioma (LGG). We corrected Spearman and Cox p-values for multiple testing using the Holm-Bonferroni method^[63]^.

## Abbreviations

(ACC): Adrenocortical carcinoma,
(BLCA): Bladder Urothelial Carcinoma,
(BRCA): Breast invasive carcinoma,
(CESC): Cervical squamous cell carcinoma and endocervical adenocarcinoma,
(CHOL): Cholangiocarcinoma,
(CLL): Chronic lymphocytic leukemia,
(COAD): Colon adenocarcinoma,
(COCOA): Coordinate Covariation Analysis,
(DLBC): Lymphoid Neoplasm Diffuse Large B-cell Lymphoma,
(ER): Estrogen receptor,
(ESCA): Esophageal carcinoma,
(FCS): Feature contribution score,
(FDR): False discovery rate,
(GBM): Glioblastoma multiforme,
(HNSC): Head and Neck squamous cell carcinoma,
(KICH): Kidney Chromophobe,
(KIRC): Kidney renal clear cell carcinoma,
(KIRP): Kidney renal papillary cell carcinoma,
(LAML): Acute Myeloid Leukemia,
(LF): Latent factor,
(LIHC): Liver hepatocellular carcinoma,
(LUAD): Lung adenocarcinoma,
(LUSC): Lung squamous cell carcinoma,
(MESO): Mesothelioma,
(MOFA): Multi-omics factor analysis,
(OV): Ovarian serous cystadenocarcinoma,
(PAAD): Pancreatic adenocarcinoma,
(PCA): Principal component analysis,
(PCPG): Pheochromocytoma and Paraganglioma,
(PRAD): Prostate adenocarcinoma,
(REA): Repressor of estrogen activity,
(READ): Rectum adenocarcinoma,
(SARC): Sarcoma,
(SKCM): Skin Cutaneous Melanoma,
(STAD): Stomach adenocarcinoma,
(TCGA): The Cancer Genome Atlas,
(TGCT): Testicular Germ Cell Tumors,
(THCA): Thyroid carcinoma,
(THYM): Thymoma,
(UCEC): Uterine Corpus Endometrial Carcinoma,
(UCS): Uterine Carcinosarcoma,
(UVM): Uveal Melanoma

## Declarations

### Ethics approval and consent to participate

Not applicable.

### Authors’ contributions

JTL led the software development with contributions from JPS and NCS. JTL, FEG-B, and NCS contributed to the writing of the manuscript. FEG-B, SB, and NCS contributed technical expertise. All authors approved the final manuscript.

## Acknowledgements

The results published here are in part based upon data generated by the TCGA Research Network: https://www.cancer.gov/tcga.

## Availability of Data and Materials

Information on the source of public data can be found in the corresponding Methods section. The R scripts used for this analysis can be accessed at https://github.com/databio/COCOA_paper. The COCOA package can be accessed at http://bioconductor.org/packages/COCOA.

## Supplementary file 1: Supplementary Methods and Information

### The power of covariation in analysis of epigenetic heterogeneity

Covariation of the epigenetic signal in different regions is an important principle in epigenetic analysis but is not fully taken advantage of by many epigenetic analysis methods. There are two common limitations of analysis methods. First, relying on differential signals between discrete sample groups loses information about the differences among samples within a group. For instance, in a health-related differential analysis, patients in the “disease” group are considered equal for the analysis when there may actually be differences between patients in the severity of their disease. Although some variation can be effectively summarized by discrete groups, in some cases, it is often more appropriate to consider variation along a continuous spectrum^[64]^. Using a continuous spectrum for samples based on physical or molecular phenotype instead of discrete groups could provide greater resolution for identifying epigenetic features that covary with sample status. Second, the use of discrete cutoffs for identifying significant epigenetic differences between samples loses information about the strength of covariation between epigenetic features and sample status. For example, epigenetic differences between samples are often determined using a discrete threshold that places epigenetic features into two groups - significantly different or not significantly different - as is done when finding differentially methylated or differentially accessible regions. Then the significant regions can be annotated with reference region sets through region set enrichment testing to aid interpretation^[1,8,65–68]^. However, while this is a flexible approach, converting continuous epigenetic signals to a binary classification -- significant or not significant -- results in the loss of covariation information that could be valuable for the region set enrichment analysis. This choice is a trade-off between the computational efficiency that comes from a simplified representation of the epigenetic signal and the potential gains that could come from having higher resolution data and most region set enrichment tools choose the simpler approach.

### Selection of immune cell-specific ATAC-seq region sets

We retrieved an ATAC-seq count matrix (GSE74912_ATACseq_All_Counts.txt.gz) from Gene Expression Omnibus with hematopoietic ATAC-seq data from Corces et al.^[69]^. We normalized each sample with quantile normalization first then GC normalization with the cqn R package^[70]^, according to the normalization done by Corces et al.^[69]^. When there were multiple samples of a given cell type from a single individual, we calculated the mean of each region to combine them into a consensus count vector. From the counts for a given cell type from various individuals, we calculated the mean in each region to create a consensus set of counts for that cell type. To get custom hematopoietic region sets, we did a series of comparisons between cell type count profiles to determine regions that were open in one or a specific group of cell types and closed in another cell type or group of cell types. We counted regions as specific when they were in the top 10% of regions in the chosen cell type/s and in the bottom 50% of the other compared cell type/s. The code for creating these region sets is available in the 0- ClusterHemaATAC.R file.

### Creation of simulated data

To create simulated data to test COCOA, we first calculated an aggregate healthy DNA methylation profile by averaging the DNA methylation profiles of 160 TCGA healthy kidney samples. To get a true positive region set, we selected an arbitrary region set (ER) and set the DNA methylation of all CpGs that overlapped that region set to zero in the healthy sample. For our analysis, we used 10 replicates of the healthy sample. We created 10 artificial disease samples by changing the DNA methylation of the CpGs in the region set of interest to between 0.0125 and 0.25, depending on the sample, with all CpGs in a given sample being assigned the same DNA methylation level. This results in covariation of the DNA methylation level of CpGs in the region of interest across samples and in differential methylation between healthy and disease samples. Finally, we added Gaussian noise to each CpG for each sample to create variation between samples, keeping methylation in the 0-1 range. We created two sample sets with different noise levels: low noise (mu=0, sd=0.025) and high noise (mu=0, sd=0.05).

To create region sets with a range of p-values, we made a set of region sets that had varied proportions of true positive regions and random loci sampled from the simulated data DNA methylation coordinates. Each random locus was expanded from the center to be 500 bp. To assign p-values to the region sets, we performed PCA on the high noise simulated data then ran COCOA on PC1 and PC2 with our region sets as the region set database. We calculated 100,000 permutations to determine empirical p-values for each region set. For further analysis in gamma approximation simulations, we selected region sets with empirical p-values across a range of orders of magnitude.

### How to choose a method for quantifying variation

The choice of method for quantifying variation depends on the data and how well that method can prioritize features that covary with each other or with a sample phenotype of interest. The decision to use covariation or correlation depends on whether the epigenetic data is proportion-based, such as for bisulfite sequencing, or count-based and unbounded, such as for ATAC-seq. This decision is not expected to greatly affect the analysis but using correlation might give greater weight to epigenetic features with very small absolute changes across samples that actually represent noise and not real signal. Using covariation may be better for proportion-based data, such as for bisulfite sequencing, and correlation may be better for count-based data, such as for ATAC-seq. Since the concept of COCOA is based on the covariation of epigenetic features across samples, COCOA will work best with methods that prioritize covarying/correlated features and do not give lower scores or coefficients to correlated features. For instance, a simple regression gives coefficients to input variables based on their association with a dependent variable. However, if two input variables are correlated, regression will give a lower coefficient to one of two variables. PCA, on the other hand, can give a high loading value to both correlated variables. An assumption of our method is that a single regulatory signal will be related to multiple regions that are regulated in a coordinated way and therefore covary across samples. For example, we would expect that the epigenetic signal in cell type-specific regions would covary across samples depending on how much of each sample corresponded to that cell type. Therefore, we expect that COCOA would work best with methods that do not lower the coefficients or scores of variables that covary. While we generally used linear metrics for quantifying variation in this study, we expect that nonlinear metrics such as feature importance scores from machine learning models would also work for quantifying epigenetic variation if they meet the criteria described above.

Some readers may notice that we use covariation or correlation instead of simply using the PCA loadings as feature contribution scores for unsupervised COCOA. Since the principal component loadings also represent the contribution of each feature to the respective principal component, we could have used those as the feature scores. However, this would have required us to recompute the PCA for each COCOA permutation to get new loadings. Instead, for each permutation, we shuffle the PC scores and calculate the covariance or correlation between the shuffled PC scores and the epigenetic data. This allows us to get new feature scores for each permutation without recalculating the PCA for every permutation, which would be computationally expensive.

### Gamma distribution p-value approximation

We used simulated data to compare empirical p-values from COCOA to the p-values derived from a gamma approximation. As mentioned earlier, we created simulated DNA methylation data with variation in the regions of a specific region set. We also created a collection of region sets that had varied similarity to the true positive region set and calculated their empirical p-values with 100,000 permutations of COCOA. To evaluate the accuracy of the gamma distribution approximation, we subsampled from the 100,000 permutations and used the subsampled COCOA runs to create gamma p-values. We did this for three subsample sizes: 300,1000, and 10,000. For each subsample size, we sampled 500,000 times, calculating the gamma p-values each time. As seen in Fig. S12, the median gamma p-value is fairly close for high p-values but tends to be lower than the empirical p-values as the p-value decreases. Increasing the number of permutations from 300 to 10,000 reduced the variance of the gamma p-values but did not cause them to converge to the empirical p-values (Fig. S12). Because of this, we recommend caution when interpreting gamma p-values, with the reminder that it is an approximation. The main benefit of the gamma p-value approximation is to screen out region sets that are not significant, which are the region sets whose p-values fall in the range where the gamma p-value approximation is more accurate.

### Considerations when choosing a region set database

The choice of region set database depends partially on the goals of the analysis but a broad database with region sets from a variety of transcription factors and cell types should be sufficient for most exploratory analyses. Along those lines, the region sets we used from ENCODE, Roadmap Epigenomics, and other sources provide a reasonably broad sampling of transcription factors and histone modification regions for a variety of cell lines and tissue types. However, any similar source of region sets could be used. The curation of region set databases is an active research area. Additionally, new region sets are continually being made available to the public. The user would benefit from any source of region sets that is relevant to their experimental question. This includes region sets derived from a cell or tissue type that is similar to the samples being studied, especially because transcription factor binding and many epigenetic marks including DNA methylation can be cell type-specific. If the user is asking a very targeted question about a specific transcription factor or cell type, the user may want to find a published region set through a source such as the Gene Expression Omnibus and use that region set alongside a broader region set database. While the database we used is not comprehensive, it is a rich starting point that can be expanded in the future.

### Other COCOA parameters

#### Absolute value of FCS

After generating the feature contribution scores (FCS), the COCOA user has the option of taking the absolute value of those scores before scoring the region sets. This choice depends on whether all regions in a region set are expected to be regulated in the same way or not (i.e. all regions activated/all regions repressed or some regions activated and some regions repressed). For cases where regions in a region set are regulated in the same direction (all activated or all repressed), it would be better to not take the absolute value. Since the FCS for important regions should all have the same sign in this case, the relevant signal will be preserved during the COCOA aggregation step while the noise from irrelevant epigenetic features, which should have arbitrary FCS signs, will cancel out. For example, a TF might activate all regions where it binds and we would expect that the epigenetic signal in these regions would generally change in the same direction and have FCS with the same sign. For cases where regions in a region set are regulated in opposite directions (some activated and some repressed), the absolute value should be taken. Since the relevant signal may have some positive and some negative FCS, aggregating FCS without taking the absolute value would partially cancel out and diminish the signal. For example, a TF might activate some regions but repress others depending on what other proteins are binding with it at a given region. In this case, the epigenetic signal in regions bound by the TF might change in opposite directions, leading to FCS with opposite signs. When taking the absolute value, it is still possible to identify region sets where regions all change in the same direction. However, FCS that represent noise will not cancel out, potentially reducing the ability to discriminate between true signal and noise. In this study, we took the absolute value of the FCS when running COCOA since there may have been some region sets in our database with regions that are regulated in opposite directions.

#### Scoring based on mean versus median

COCOA offers the option to score based on the median region set FCS instead of the mean FCS. To compare the median scoring method to the mean scoring method which was used in the main text, we performed COCOA with the median scoring method on the TCGA breast cancer DNA methylation data. We see that the overall trends are similar, with ER-related region sets found to be highly ranked for PC1 and PC3 and polycomb-related region sets highly ranked for PC4 (Fig. S11A, Additional File 1: Table S14). Additionally, the meta-region profiles for top region sets from the mean scoring method also have peaks for the median scoring method (Fig. S11B). Consistent with these observations, the region set scores for the first 4 PCs have very high Spearman correlation between scoring methods, all with at least 0.95 correlation (Fig. S11C).

### Discussion of EZH2 results in comparison to previous findings

Several trends present in our EZH2/SUZ12-binding region analysis contrast with previous results. First, we found a significant positive correlation between EZH2-binding region DNA methylation and cancer stage in testicular germ cell tumors (TGCT), whereas previous studies did not identify an association between EZH2 expression and cancer stage^[71]^ and suggested that EZH2 activity is decreased during cancer progression^[71]^ and in chemotherapy resistance^[72]^. Second, we found a negative correlation between EZH2-binding region methylation and cancer stage in UVM that trended toward significance (uncorrected p<0.05) while a previous study suggested that increased expression of EZH2 was positively associated with higher risk of metastasis^[30]^. Third, our finding that higher DNA methylation in EZH2- binding regions trended toward significance (uncorrected p<0.05) for association with lower risk of death in GBM contrasts with reports suggesting that EZH2 expression promotes proliferation and tumorigenesis in glioblastoma^[73,74]^. These trends could be due in part to the context-dependent effects of EZH2^[31,36,37]^. Further studies would be valuable to clarify the role of EZH2 in these cancer types.

### Comparison of COCOA to other region set or covariation-based methods

We are not aware of any other tool designed for DNA methylation data that identifies region sets based on DNA methylation variation across samples. However, since COCOA is broadly applicable to epigenetic data, we provide a comparison between COCOA and tools designed for chromatin accessibility data with which COCOA shares some important concepts. We also compare COCOA to tools that were not designed for epigenetic data but have some conceptual similarity to COCOA. Finally, we mention a tool designed for DNA methylation data that has superficial similarity to COCOA but actually performs a very different function. COCOA is unique in that it provides a class of DNA methylation heterogeneity analysis that was not previously available but also provides a framework to apply the same method to other epigenetic data types.

#### Tools for chromatin accessibility data

##### ChromVAR

ChromVAR is an R package that quantifies the variability of chromatin accessibility signal in motif regions or region sets^[2]^. For a given set of motif regions, each sample is given a score for how much it deviates from the expected chromatin accessibility of those motif regions. Each motif region set is also given a score for how variable it is across samples. ChromVAR has a few major differences from COCOA. First, as mentioned previously, COCOA works with DNA methylation data or chromatin accessibility data, while chromVAR was designed for chromatin accessibility data. Second, COCOA can use multiple metrics to quantify epigenetic variation across samples while chromVAR only uses a single unsupervised way of quantifying variation (bias-corrected z-score for each sample, region set combination). Among COCOA’s multiple options, COCOA can use PCA to more easily separate and annotate biological signals. COCOA also supports supervised analysis, adding the ability to do a range of new analyses not supported by chromVAR. Third, a smaller point, COCOA includes additional data analysis and visualization functions such as for meta-region profiles to further understand inter-sample variation. While chromVAR’s utility is attested to by the many papers citing it, COCOA adds meaningful value to the epigenetics field that is not captured by the chromVAR package.

##### BROCKMAN

BROCKMAN is a tool designed primarily for single cell chromatin accessibility data that uses variation in the frequency of k-mers in reads to identify gene regulatory variation across cells^[9]^. While BROCKMAN and COCOA share some conceptual foundations, specifically that covariation of regulatory signals across cells or samples can be used to understand gene regulatory differences between the cells, there are some important differences. First, the BROCKMAN tool is for chromatin accessibility data, not DNA methylation, while COCOA has a generalized framework that works for both data types. Second, BROCKMAN aggregates epigenetic signal by category (k-mer) before doing dimensionality reduction while COCOA first does dimensionality reduction (or other quantification method) then aggregates epigenetic signal by category (region set). Aggregating before dimensionality reduction is well suited to single cell data, as has been done for single cell DNA methylation data ^[75]^. However, aggregating epigenetic signal after dimensionality reduction allows more flexibility in applications and allows genome-wide variability to be captured in a more unbiased way. For example, COCOA could be used with multi-omic dimensionality reduction as shown in Figure 4 with minimal changes to the COCOA algorithm. Aggregating signal within region sets first might miss inter-sample epigenetic variability that is not contained within any tested region sets. COCOA shares some ideas with BROCKMAN but applies them in a generalized framework that can apply to new epigenetic data types, including DNA methylation.

#### Gene-centric methods with conceptual similarity to COCOA

The next three methods have some conceptual overlap with COCOA but are gene-centric rather than region-centric. As mentioned in the paper introduction, region-based approaches are more appropriate for epigenetic data, for reasons including that it can be difficult to link epigenetic marks to genes.

##### PCGSE

Principal component gene set enrichment (PCGSE) is a method to annotate principal components that are derived from gene expression data with gene sets^[10]^. COCOA derived conceptual foundations from this method but extends them to apply to epigenetic data and region sets. COCOA also extends beyond PCA to include other analyses including supervised analysis.

##### MOGSA

Multi-omics gene set analysis (MOGSA) uses matrix factorization on multi-omics data from the same samples to integrate the data and reduce its dimensionality then does gene set analysis^[12]^. This method is gene-centric and not tailored to epigenetic data. As shown with MOFA in Fig. 4, multi-omics dimensionality reduction techniques could benefit from including a region-centric method such as COCOA to annotate the epigenetic component of inter-sample variation in addition to using gene set analysis

##### PathwayPCA

PathwayPCA can do pathway analysis in a variety of scenarios using supervised PCA and Adaptive Elastic-net Sparse PCA^[13]^. This method is gene-centric and is focused on pathways. As such, it has a different focus than COCOA.

#### Method for DNA methylation that uses local covariation

##### CoMethDMR

CoMethDMR is a tool to identify differentially methylated regions (DMRs)^[76]^. To boost statistical power, CoMethDMR takes into account local covariation of DNA methylation within a given region. Unlike CoMethDMR which uses covariation of the epigenetic signal only locally, COCOA uses covariation of the epigenetic signal on the genome-scale. While CoMethDMR and COCOA may have superficial similarities, their goals are different. The output of CoMethDMR is a set of differentially methylated regions while the output of COCOA is a list of region sets associated with a target variable.

### Comparison of COCOA to ChromVAR

We compared COCOA and chromVAR with two main comparisons with the breast cancer ATAC-seq data: both tools applied with the main database of region sets used for this paper and both tools applied with the curated motif database used in the chromVAR paper. For the first comparison, both methods rank ER and ER-related region sets highly although COCOA did this to a greater extent (Fig. S9A), perhaps because the use of PCA for COCOA allowed it to separate epigenetic signals more clearly. The median rank for ER region sets was 45 for PC1 of COCOA and 607 for chromVAR, with 31 ER region sets in the database. ChromVAR also did not rank hematopoietic transcription factors highly, as PC2 of COCOA did (Fig. S9A), but many of the highest scoring region sets for chromVAR were region sets for histone modifications or chromatin accessibility in immune cells. It is possible that COCOA and chromVAR are uncovering the same underlying signal but in different ways. For the second comparison, ER motifs were not ranked highly for PC1 of COCOA or for chromVAR (Fig. S9B), which may be due to differences between ER motif regions and ER ChIP-seq data, although chromVAR ranked ER motifs higher than COCOA. The median rank for ER motifs was 1309 for PC1 of COCOA and 216 for chromVAR, with 3 ER motifs in the database. Both chromVAR and PC1 of COCOA identified FOXA1 as the highest scoring ER- related motif (Fig. S9B). Both chromVAR and COCOA also identified many other FOX motifs as top results (Fig. S9C), presumably because of their similarity to FOXA1. Some of chromVAR’s highest scoring motifs were for AP1 components. AP1 colocalizes with ER and may be a tethering factor for ER^[77,78]^. While PC1 of COCOA does not rank AP1-related motifs highly, PCs 3 and 4 do rank them highly (Fig. S9C). COCOA did not rank motifs for hematopoietic TFs highly for PC2 as it did for the region sets for hematopoietic TFs, although some hematopoietic TF motifs do have high scores for PC4 (Fig. S9C), which is more consistent with the region set results. This may once again be due to the difference between motifs and ChIP-seq region sets.

We observe that both COCOA and chromVAR achieved higher maximum scores for experimental region sets (Fig. S9A) than for motifs (Fig. S9B, S9C) although this trend could depend on the cutoff for determining motif matches (default parameters were used). For instance, the COCOA score (average absolute correlation) for PC1 for the highest ranking and median ER region sets were 0.52 and 0.40 while the scores for the highest ranking and median ER motifs were both 0.33. The chromVAR scores (standard deviation of samples’ z-scores) for the highest ranking and median ER region sets were 47.11 and 28.14 while the scores for the highest ranking and median ER motifs were 14.23 and 13.52. Because the region set database performed better for both methods, we argue that the results using the region set database are more relevant for comparing the methods. Overall, our results demonstrate that both methods can discover relevant biological insights but COCOA can separate biological signals to a greater extent than chromVAR since COCOA’s flexible framework allows the use of PCA.

### Comparison of COCOA to LOLA

We compared COCOA to a generic region set enrichment method that does not consider covariation, LOLA^[1]^, which is a previous method associated with our lab. We performed two comparisons of COCOA and LOLA with simulated data: one in which we added a low level of noise to our samples and one in which we added a higher level of noise (Fig. S9A). Since LOLA requires a set of regions as input, we used the bumphunter R package^[79]^ to find differentially methylated regions (DMRs) between healthy and disease samples. Then, we used LOLA to test the DMRs for enrichment against our region set database. For COCOA, we performed PCA on the simulated samples then identified region sets associated with PC1 and PC2. For the comparison with a low level of noise, both methods were able to identify the region set of interest (Fig. S9B, Additional file 1: Tables S10, S11). However, with a higher level of noise, bumphunter did not identify any significant DMRs (FDR < 0.05) and we were therefore unable to run LOLA (Fig. S9C). In contrast, COCOA was still able to identify the region set of interest as relevant for PC2 (Fig. S9C, Additional file 1: Table S12). In this case, the noise apparently begins to dominate the variation among samples, and noise is therefore detected in PC1. However, the signal is still present, and is now detected by COCOA in PC2. Despite the noise, COCOA can still discover the healthy vs disease signal as relevant in PC2, while the bumphunter + LOLA approach is not able to detect significant differences. This comparison demonstrates that COCOA can better leverage the covariation of epigenetic signal to annotate epigenetic variation compared to methods that do not use covariation.

**Fig. S1.**
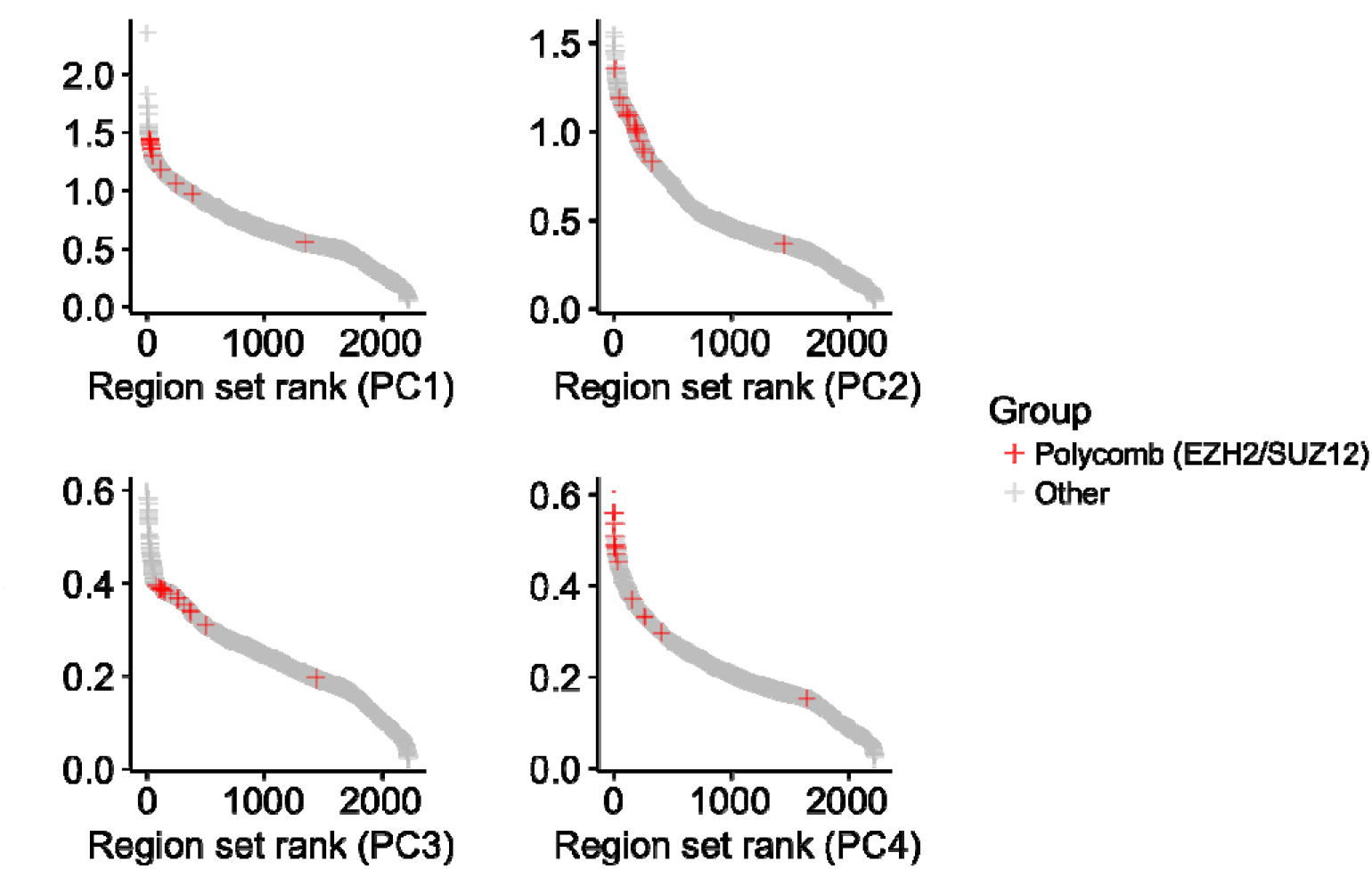
Region set scores for PCs 1-4 for the BRCA DNA methylation data. This figure is included with only the polycomb group marked to allow clearer visualization of the polycomb region set group in comparison to Fig. 1 where several region set groups are marked.

**Fig. S2.**
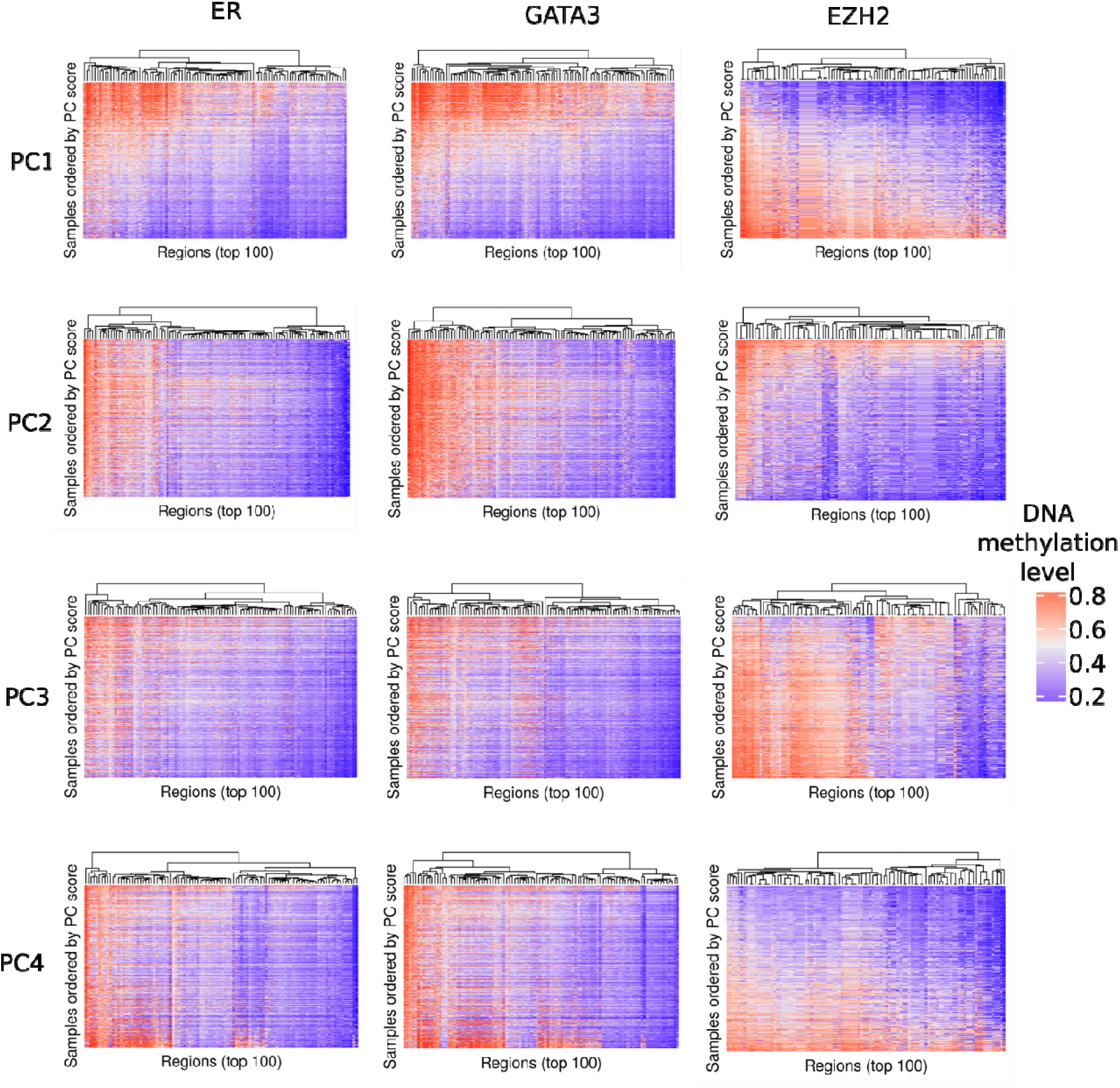
DNA methylation in some of the top scoring region sets for principal component 1. Average DNA methylation levels are shown for the 100 regions from each region set that had the highest absolute FCS for each PC. Patients are ordered by PC scores.

**Fig. S3.**
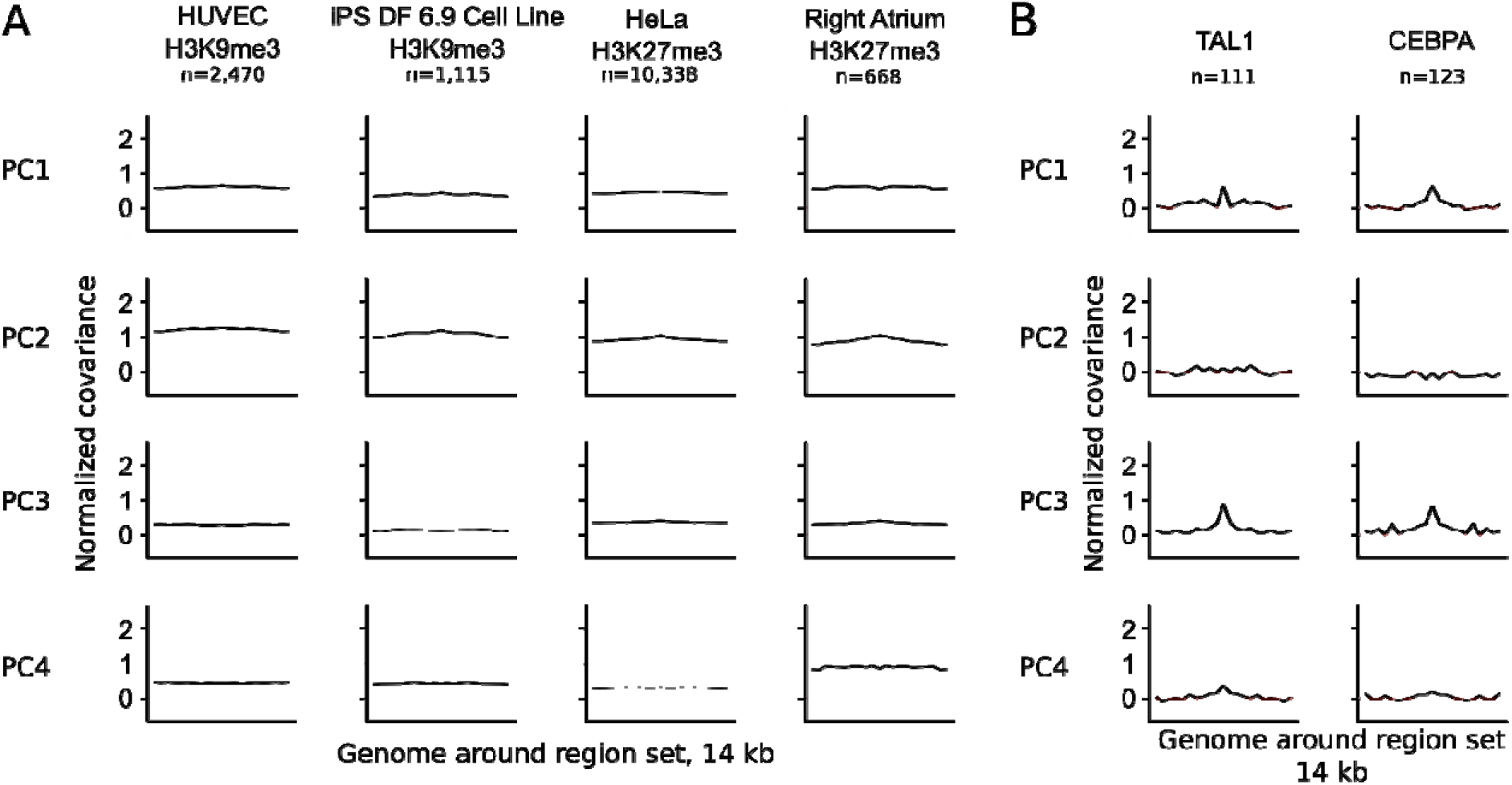
Meta-region profiles for region sets from the COCOA analysis of breast cancer DNA methylation data. A. Profiles for the highest scoring H3K9me3 and H3K27me3 region sets from PC2. B. Profiles for the two highest scoring hematopoietic TFs in PC3. A peak in the center of the meta-region profile indicates that the DNA methylation level covaries with the PC more at the region of interest than in the surrounding genome. Profiles have been normalized to the mean and standard deviation of the covariance of all cytosines for each PC. The number of regions from each region set that were covered by the epigenetic data in the COCOA analysis (Fig. 2, panel A) is indicated by “n”.

**Fig. S4.**
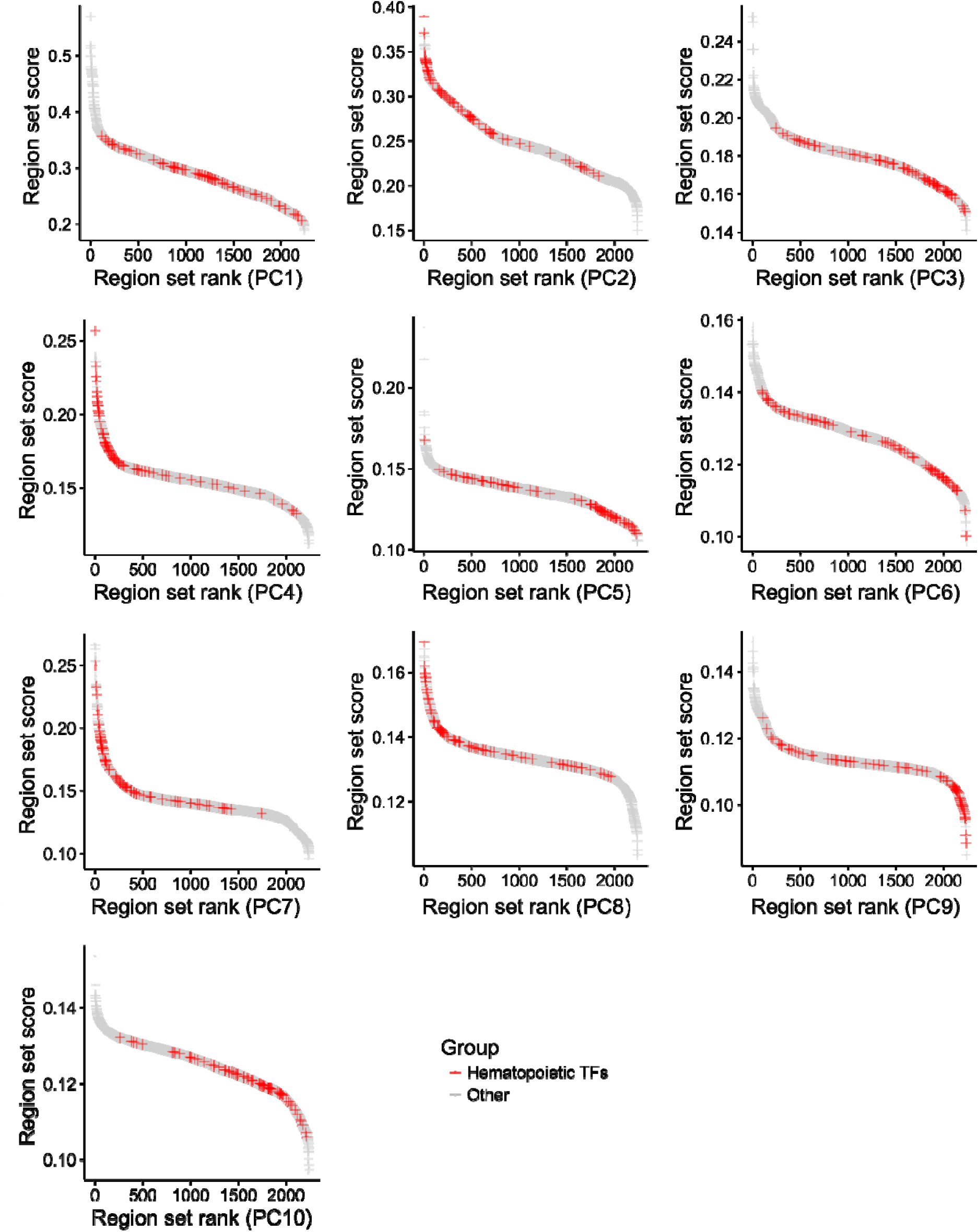
Hematopoietic transcription factor region sets have high scores for several of the top principal components. Region set scores for each of the first 10 principal components of the BRCA ATAC-seq data.

**Fig. S5.**
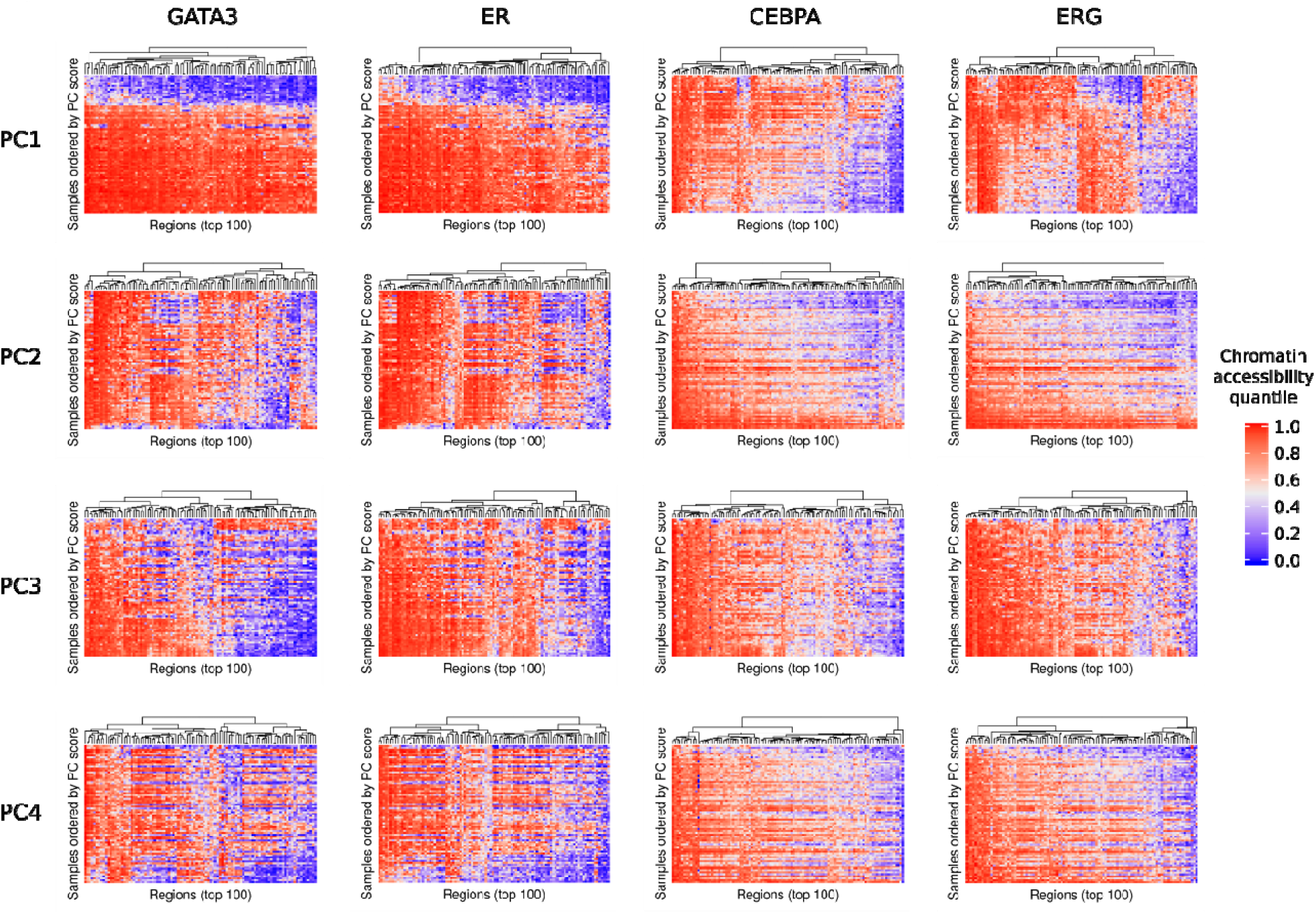
Chromatin accessibility signal in some of the top scoring region sets from COCOA analysis of breast cancer ATAC-seq data. GATA3 and ER were the top scoring region sets for PC1 while CEBPA and ERG were the top scoring region sets for PC2. Average chromatin accessibility quantiles are shown for the 100 regions from each region set that had the highest absolute FCS for each PC. Patients are ordered by PC scores.

**Fig. S6.**
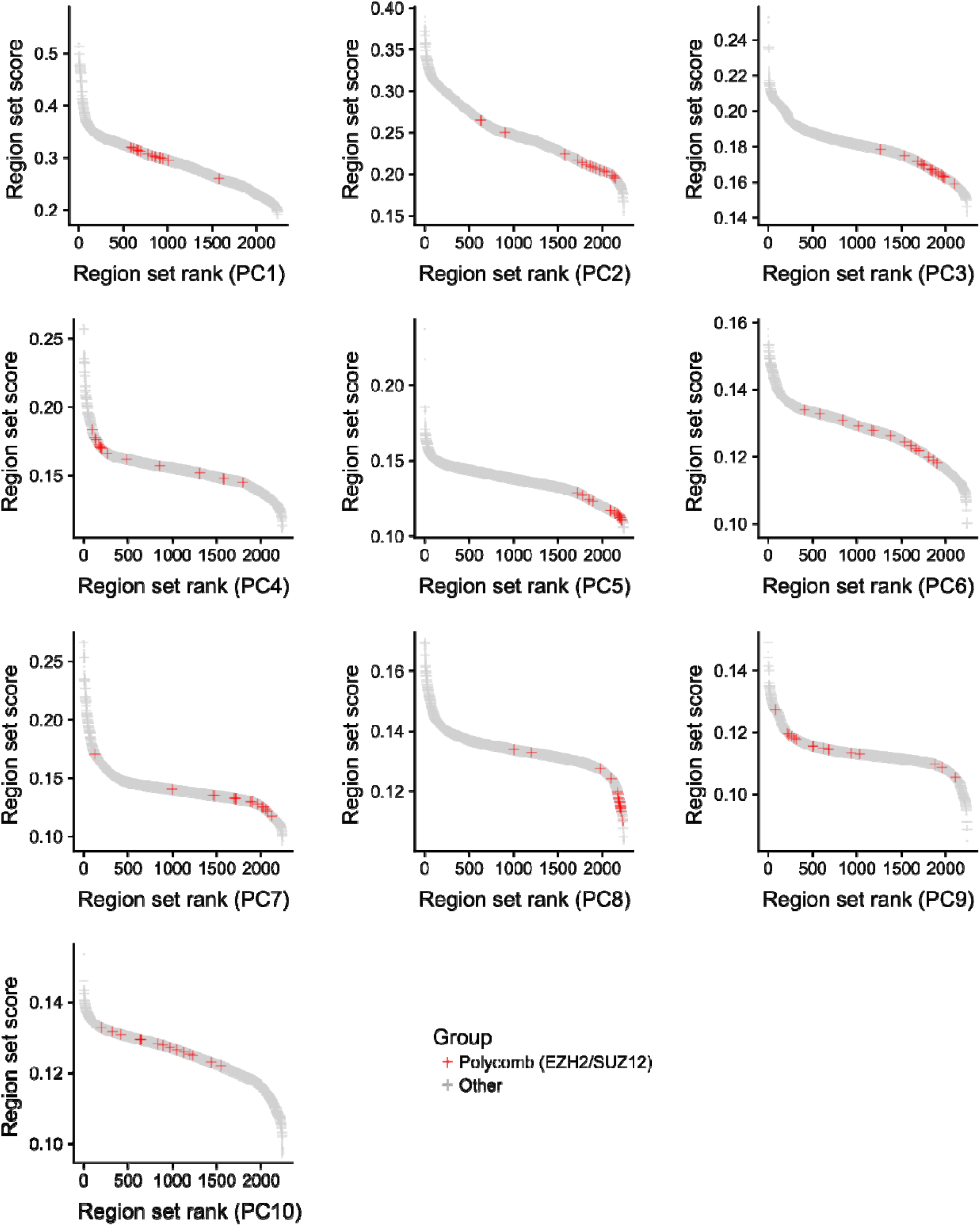
Region set scores for each of the first 10 principal components of the BRCA ATAC-seq data, with polycomb region sets (EZH2/SUZ12-binding regions) indicated.

**Fig. S7.**
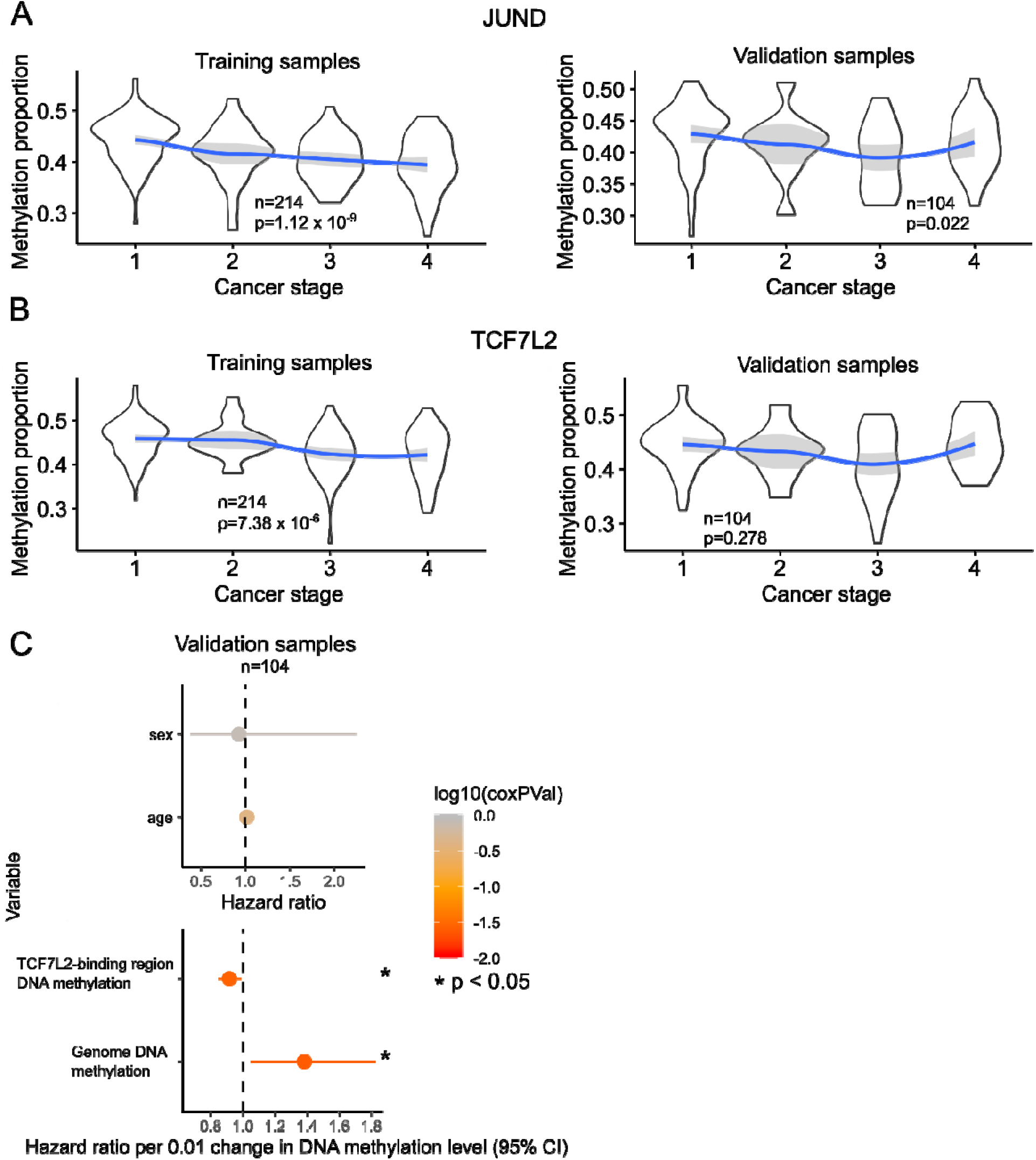
Association of average DNA methylation level in JUND and TCF7L2-binding regions with KIRC cancer stage and overall survival. A. The Spearman correlation of cancer stage with the average DNA methylation in JUND-binding regions. The JUND region set used is the highest scoring transcription factor region set from the KIRC COCOA analysis. The JUND Cox proportional hazards model did not meet the proportional hazards assumption and is therefore not included in the figure. B. The Spearman correlation of cancer stage with the average DNA methylation in TCF7L2-binding regions. The TCF7L2 region set used is the second highest scoring transcription factor region set from the KIRC COCOA analysis. C. Hazard ratios for Cox proportional hazards model of the association between overall patient survival and average DNA methylation in TCF7L2-binding regions.

**Fig. S8.**
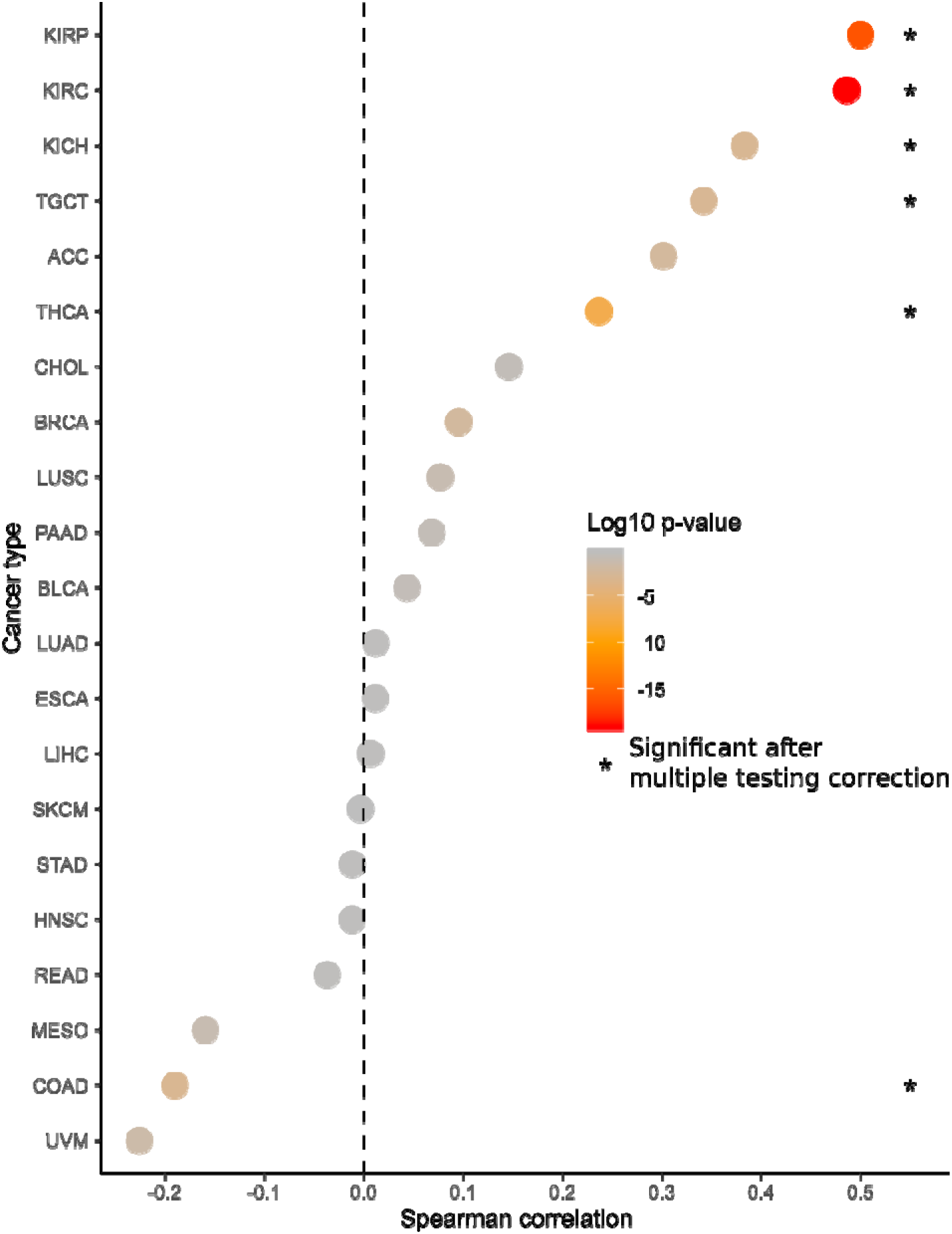
Correlation between average EZH2/SUZ12-binding region DNA methylation and cancer stage. Color is based on the raw Spearman p-values and asterisks mark significant correlations after Holm-Bonferroni correction to account for testing 21 cancer types.

**Figure S9.**
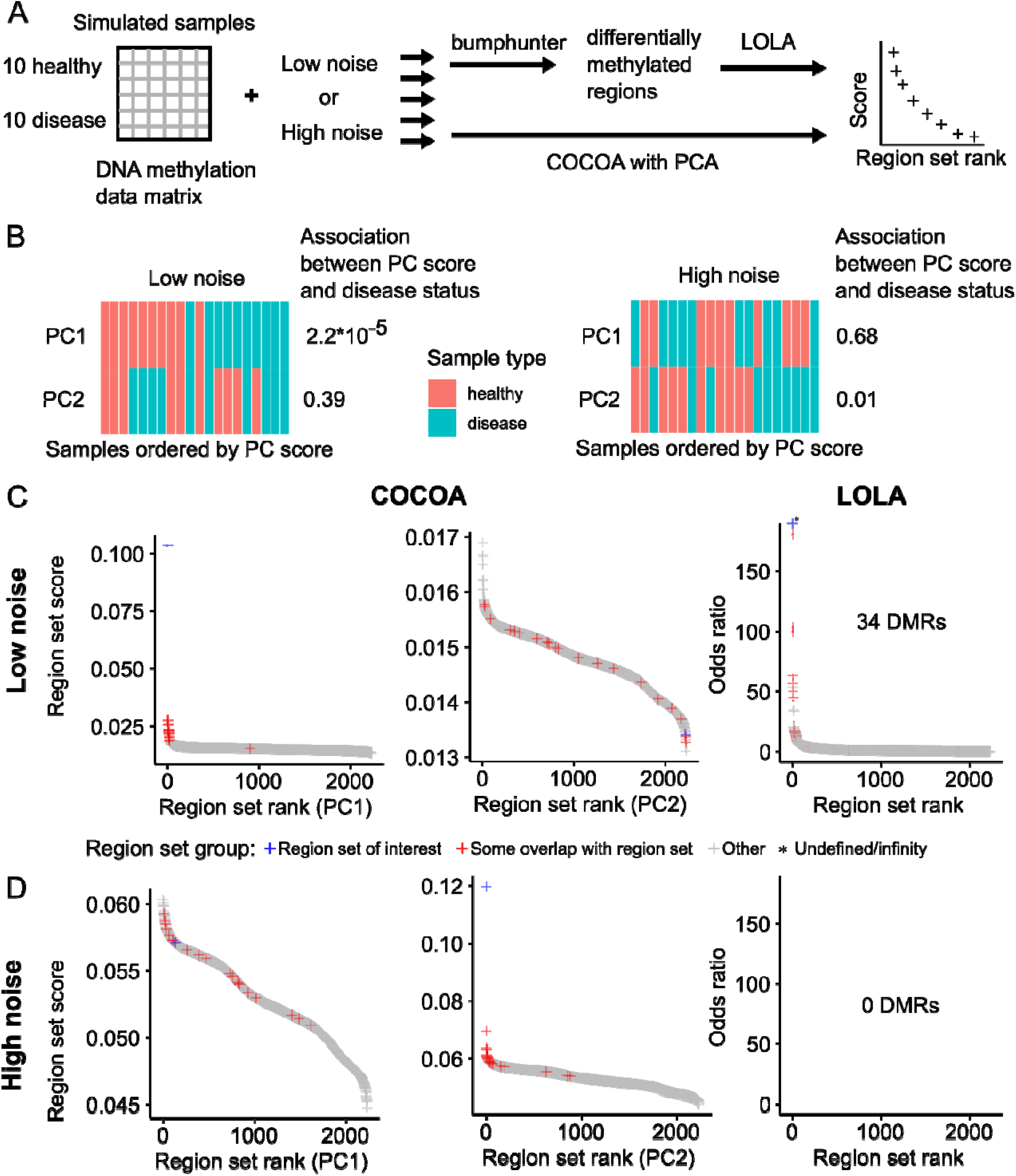
Comparison of COCOA and LOLA. A. The workflow for comparison of the methods. B. Association of PC scores with disease status. For low noise, PC1 is associated with disease status but for high noise, PC2 is associated with disease status (Wilcoxon rank-sum test). C. Results with a low level of noise added to samples. The COCOA score or LOLA odds ratio for each region set, ordered from highest to lowest. C. Results with a high level of noise added to samples. The COCOA score or LOLA odds ratio for each region set, ordered from highest to lowest. There are no scores for LOLA because bumphunter did not identify any significant DMRs (FDR <= 0.05).

**Fig. S10.**
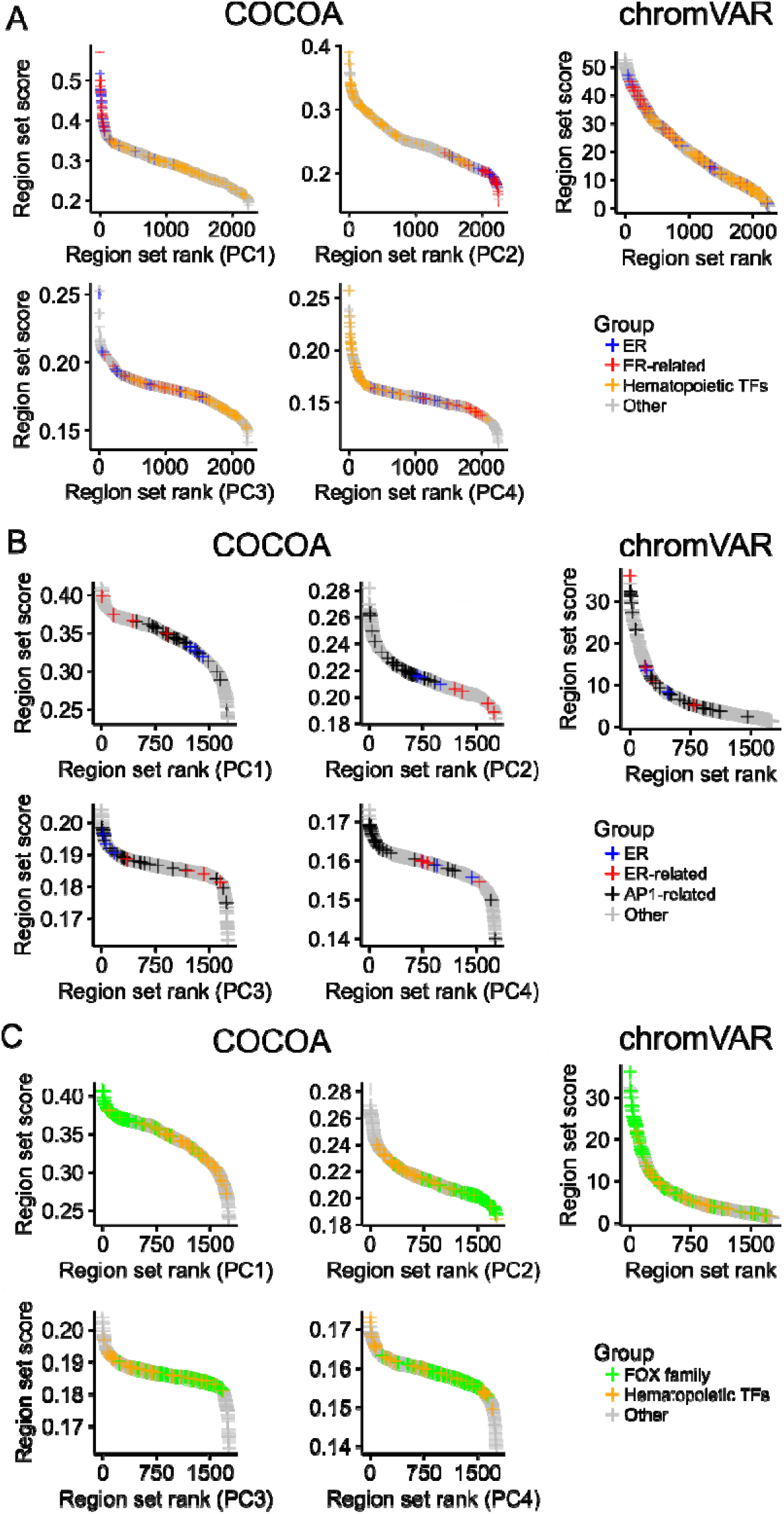
Comparison of COCOA and chromVAR on breast cancer ATAC-seq data. A. COCOA and chromVAR scores for the region set database (see “Region set database” in methods). The chromVAR score for a region set is the standard deviation of all samples’ chromatin accessibility z-scores for that region set. The ER-related region set group includes FOXA1, GATA3, and H3R17me2. For definition of the hematopoietic TF group, see “Region set database” in methods. B. COCOA and chromVAR scores for a curated version of the cisBP motif database. The ER-related region set group includes FOXA1 and GATA3. For the definition of the AP1-related group, see “Comparison of COCOA and chromVAR” in methods. C. The same COCOA and chromVAR scores for a curated version of the cisBP motif database but indicating FOX family motifs and hematopoietic TF motifs.

**Fig. S11.**
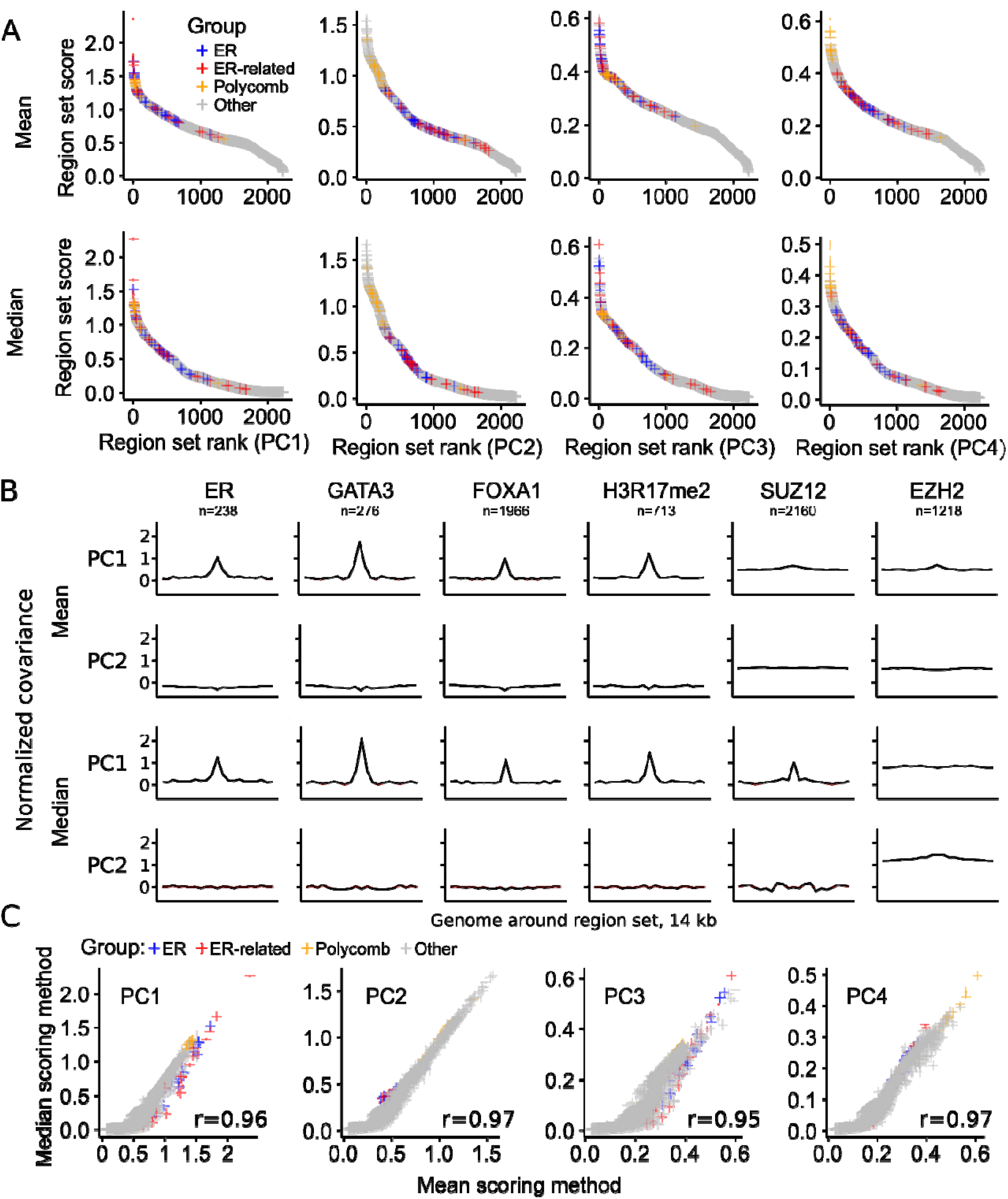
Comparison of median and mean scoring methods. A. The COCOA score for each region set, ordered from highest to lowest. The ER-related group includes GATA3, FOXA1, and H3R17me2. The polycomb group includes EZH2 and SUZ12. B. Meta-region profiles of several of the highest scoring region sets from the breast cancer analysis. Meta-region profiles show covariance between PC scores and the epigenetic signal in regions of the region set, centered on the regions of interest. The number of regions from each region set that were covered by the epigenetic data in the COCOA analysis (panel A) is indicated by “n”. The line at zero marks the mean or median respectively of the FCS for each PC. C. The relationship between region set scores for each scoring method. The Spearman correlation is shown.

**Fig. S12.**
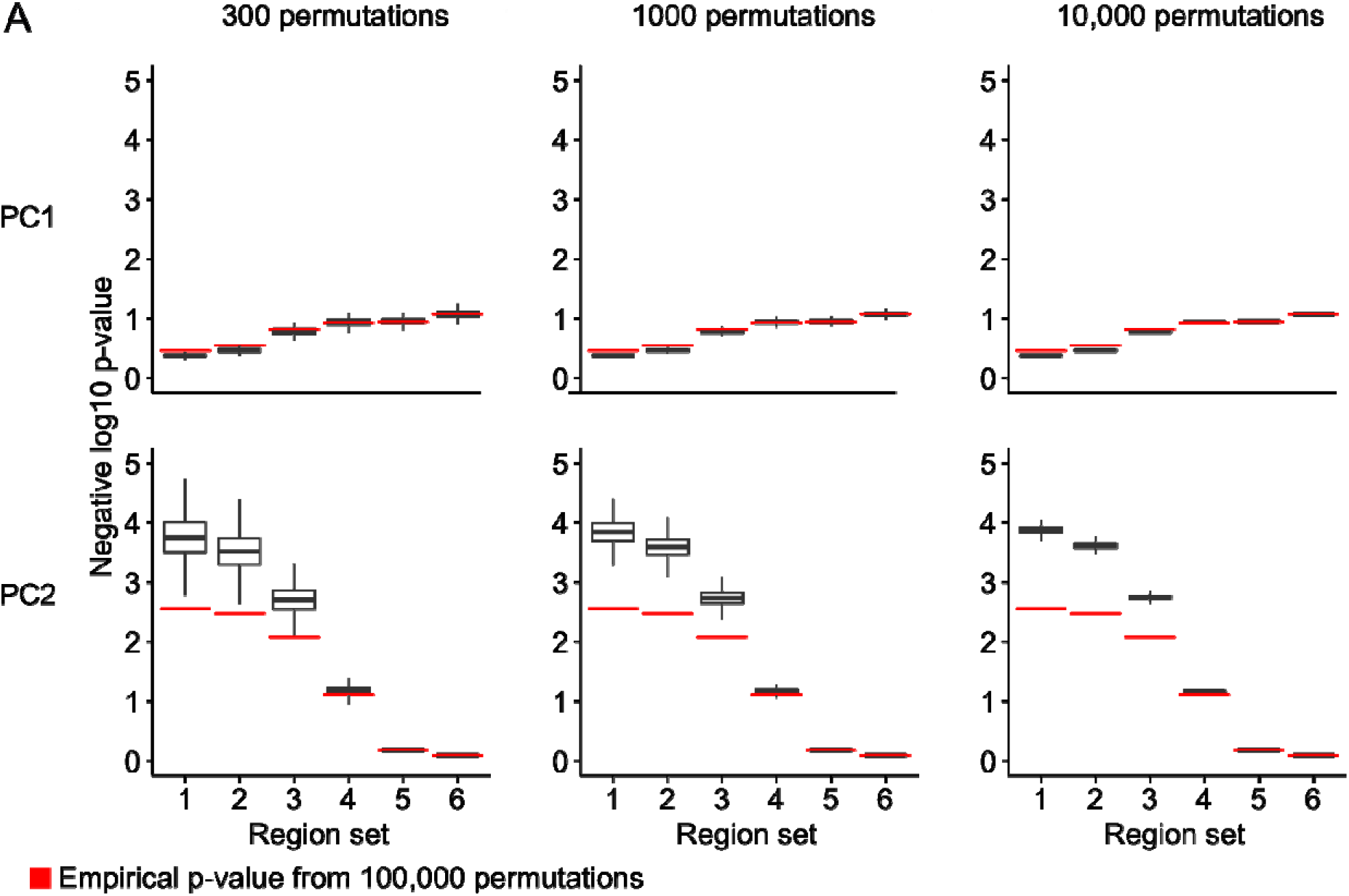
Comparison of empirical p-values to gamma distribution p-value approximation. COCOA was run on PC1 and PC2 of PCA of simulated data with six region sets. The empirical p-values from 100,000 permutations are shown. P-values were also calculated with a gamma distribution approximation after sampling either 300, 1000, or 10,000 permutations from the 100,000 that were calculated. 500,000 such samples were taken to get a distribution of gamma p-values for each region set (outliers not shown).

## References

[1] Sheffield NC, Bock C. LOLA: enrichment analysis for genomic region sets and regulatory elements in R and Bioconductor. Bioinformatics. 2015 oct;32(4):587–589.

[2] Schep AN, Wu B, Buenrostro JD, Greenleaf WJ. chromVAR: inferring transcription-factor-associated accessibility from single-cell epigenomic data. Nature Methods. 2017 aug;14(10):975–978.

[3] Lawson JT, Tomazou EM, Bock C, Sheffield NC. MIRA: an R package for DNA methylation-based inference of regulatory activity. Bioinformatics. 2018 mar;34(15):2649–2650.

[4] McLean CY, Bristor D, Hiller M, Clarke SL, Schaar BT, Lowe CB, et al. GREAT improves functional interpretation of cis-regulatory regions. Nature Biotechnology. 2010 may;28(5):495–501.

[5] Sheffield NC, Thurman RE, Song L, Safi A, Stamatoyannopoulos JA, Lenhard B, et al. Patterns of regulatory activity across diverse human cell types predict tissue identity, transcription factor binding, and long-range interactions. Genome Research. 2013 mar;23(5):777–788.

[6] Sheffield NC, Pierron G, Klughammer J, Datlinger P, Schonegger A, Schuster M, et al. DNA methylation heterogeneity defines a disease spectrum in Ewing sarcoma. Nature Medicine. 2017 jan;23(3):386–395.

[7] Dozmorov MG. Epigenomic annotation-based interpretation of genomic data: from enrichment analysis to machine learning. Bioinformatics. 2017 jun;33(20):3323–3330.

[8] Layer RM, Pedersen BS, DiSera T, Marth GT, Gertz J, Quinlan AR. GIGGLE: a search engine for large-scale integrated genome analysis. Nature Methods. 2018 jan;15(2):123–126.

[9] de Boer CG, Regev A. BROCKMAN: deciphering variance in epigenomic regulators by k-mer factorization. BMC Bioinformatics. 2018 jul;19(1).

[10] Frost HR, Li Z, Moore JH. Principal component gene set enrichment (PCGSE). BioData Mining. 2015 jun;8(1).

[11] Subramanian A, Tamayo P, Mootha VK, Mukherjee S, Ebert BL, Gillette MA, et al. Gene set enrichment analysis: A knowledge-based approach for interpreting genome-wide expression profiles. Proceedings of the National Academy of Sciences. 2005 sep;102(43):15545–15550.

[12] Meng C, Basunia A, Peters B, Gholami AM, Kuster B, Culhane AC. MOGSA: Integrative Single Sample Geneset Analysis of Multiple Omics Data. Molecular & Cellular Proteomics. 2019 jun;18(8 suppl 1):S153–Sl68.

[13] Odom GJ, Ban Y, Liu L, Sun X, Pico AR, Zhang B, et al. pathwayPCA: an R package for integrative pathway analysis with modern PCA methodology and gene selection. 2019 apr;.

[14] Ung M, Ma X, Johnson KC, Christensen BC, Cheng C. Effect of estrogen receptor alpha binding on functional DNA methylation in breast cancer. Epigenetics. 2014 jan;9(4):523–532.

[15] Fleischer T,, Tekpli X, Mathelier A, Wang S, Nebdal D, et al. DNA methylation at enhancers identifies distinct breast cancer lineages. Nature Communications. 2017 nov;8(1).

[16] Frietze S, Lupien M, Silver PA, Brown M. CARM1 Regulates Estrogen-Stimulated Breast Cancer Growth through Up-regulation of E2F1. Cancer Research. 2008 jan;68(1):301–306.

[17] Guo S, Li X, Rohr J, Wang Y, Ma S, Chen P, et al. EZH2 overexpression in different immunophenotypes of breast carcinoma and association with clinicopathologic features. Diagnostic Pathology. 2016 apr;11(1).

[18] Holm K, Grabau D, Lovgren K, Aradottir S, Gruvberger-Saal S, Howlin J, et al. Global H3K27 trimethylation and EZH2 abundance in breast tumor subtypes. Molecular Oncology. 2012 jun;6(5):494–506.

[19] Hwang C, Giri VN, Wilkinson JC, Wright CW, Wilkinson AS, Cooney KA, et al. EZH2 regulates the transcription of estrogen-responsive genes through association with REA, an estrogen receptor corepressor. Breast Cancer Research and Treatment. 2007 apr;107(2):235–242.

[20] Segovia-Mendoza M, Morales-Montor J. Immune Tumor Microenvironment in Breast Cancer and the Participation of Estrogen and Its Receptors in Cancer Physiopathology. Frontiers in Immunology. 2019 mar;10.

[21] Corces MR, Granja JM, Shams S, Louie BH, Seoane JA, Zhou W, et al. The chromatin accessibility landscape of primary human cancers. Science. 2018 oct;362(6413):eaav1898.

[22] Dietrich S, Oles M, Lu J, Sellner L, Anders S, Velten B, et al. Drug-perturbation-based stratification of blood cancer. Journal of Clinical Investigation. 2017 dec;128(1):427–445.

[23] Argelaguet R, Velten B, Arnol D, Dietrich S, Zenz T, Marioni JC, et al. Multi-Omics Factor Analysis-a framework for unsupervised integration of multi-omics data sets. Molecular Systems Biology. 2018 jun;14(6):e8124.

[24] Fabbri G, Dalla-Favera R. The molecular pathogenesis of chronic lymphocytic leukaemia. Nature Reviews Cancer. 2016 feb; 16(3): 145–162.

[25] Takao Y, Yokota T, Koide H. β-Catenin up-regulates Nanog expression through interaction with Oct-3/4 in embryonic stem cells. Biochemical and Biophysical Research Communications. 2007 feb;353(3):699–705.

[26] Faunes F, Hayward P, Descalzo SM, Chatterjee SS, Balayo T, Trott J, et al. A membrane-associated β-catenin/Oct4 complex correlates with ground-state pluripotency in mouse embryonic stem cells. Development. 2013 feb;140(6):1171–1183.

[27] Ying L, Mills JA, French DL, Gadue P. OCT4 Coordinates with WNT Signaling to Pre-pattern Chromatin at the SOX17 Locus during Human ES Cell Differentiation into Definitive Endoderm. Stem Cell Reports. 2015 oct;5(4):490–498.

[28] Zhang D, Yang X, Luo Q, Fu D, Li H, Li H, et al. EZH2 enhances the invasive capability of renal cell carcinoma cells via activation of STAT3. Molecular Medicine Reports. 2017 dec;.

[29] Varambally S, Dhanasekaran SM, Zhou M, Barrette TR, Kumar-Sinha C, Sanda MG, et al. The polycomb group protein EZH2 is involved in progression of prostate cancer. Nature. 2002 oct;419(6907):624–629.

[30] Cheng Y, Li Y, Huang X, Wei W, Qu Y. Expression of EZH2 in uveal melanomas patients and associations with prognosis. Oncotarget. 2017 jul;8(44).

[31] Kim KH, Roberts CWM. Targeting EZH2 in cancer. Nature Medicine. 2016 feb;22(2):128–134.

[32] Bachmann IM, Halvorsen OJ, Collett K, Stefansson IM, Straume O, Haukaas SA, et al. EZH2 Expression Is Associated With High Proliferation Rate and Aggressive Tumor Subgroups in Cutaneous Melanoma and Cancers of the Endometrium, Prostate, and Breast. Journal of Clinical Oncology. 2006 jan;24(2):268–273.

[33] Melling N, Thomsen E, Tsourlakis MC, Kluth M, Hube-Magg C, Minner S, et al. Overexpression of enhancer of zeste homolog 2 (EZH2) characterizes an aggressive subset of prostate cancers and predicts patient prognosis independently from pre-and postoperatively assessed clinicopathological parameters. Carcinogenesis. 2015 sep;36(11):1333–1340.

[34] Liu L, Xu Z, Zhong L, Wang H, Jiang S, Long Q, et al. Prognostic Value of EZH2 Expression and Activity in Renal Cell Carcinoma: A Prospective Study. PLoS ONE. 2013 nov;8(11):e81484.

[35] Chen Z, Yang P, Li W, He F, Wei J, Zhang T, et al. Expression of EZH2 is associated with poor outcome in colorectal cancer. Oncology Letters. 2017 dec;.

[36] Wang Y, Hou N, Cheng X, Zhang J, Tan X, Zhang C, et al. Ezh2 Acts as a Tumor Suppressor in Kras-driven Lung Adenocarcinoma. International Journal of Biological Sciences. 2017;13(5):652–659.

[37] Basheer F, Giotopoulos G, Meduri E, Yun H, Mazan M, Sasca D, et al. Contrasting requirements during disease evolution identify EZH2 as a therapeutic target in AML. The Journal of Experimental Medicine. 2019 mar;216(4):966–981.

[38] Huber W, Carey VJ, Gentleman R, Anders S, Carlson M, Carvalho BS, et al. Orchestrating high-throughput genomic analysis with Bioconductor. Nature Methods. 2015 jan;12(2):115–121.

[39] Consortium EP. An integrated encyclopedia of DNA elements in the human genome. Nature. 2012 sep;489(7414):57–74.

[40] Davis CA, Hitz BC, Sloan CA, Chan ET, Davidson JM, Gabdank I, et al. The Encyclopedia of DNA elements (ENCODE): data portal update. Nucleic Acids Research. 2017 nov;46(D1):D794–D801.

[41] Bernstein BE, Stamatoyannopoulos JA, Costello JF, Ren B, Milosavljevic A, Meissner A, et al. The NIH Roadmap Epigenomics Mapping Consortium. Nature Biotechnology. 2010 oct;28(10): 1045–1048.

[42] Kundaje A,, Meuleman W, Ernst J, Bilenky M, Yen A, et al. Integrative analysis of 111 reference human epigenomes. Nature. 2015 feb;518(7539):317–330.

[43] Winkler AM, Ridgway GR, Douaud G, Nichols TE, Smith SM. Faster permutation inference in brain imaging. NeuroImage. 2016 nov;141:502–516.

[44] Delignette-Muller ML, Dutang C. fitdistrplus: An R Package for Fitting Distributions. Journal of Statistical Software. 2015;64(4).

[45] Benjamini Y, Hochberg Y. Controlling the False Discovery Rate: A Practical and Powerful Approach to Multiple Testing. Journal of the Royal Statistical Society: Series B (Methodological). 1995 jan;57(1):289–300.

[46] Sánchez-Castillo M, Ruau D, Wilkinson AC, Ng FSL, Hannah R, Diamanti E, et al. CODEX: a next-generation sequencing experiment database for the haematopoietic and embryonic stem cell communities. Nucleic Acids Research. 2014 sep;43(D1):D1117–D1123.

[47] Mei S, Qin Q, Wu Q, Sun H, Zheng R, Zang C, et al. Cistrome Data Browser: a data portal for ChIP-Seq and chromatin accessibility data in human and mouse. Nucleic Acids Research. 2016 oct;45(D1):D658–D662.

[48] Sandelin A. JASPAR: an open-access database for eukaryotic transcription factor binding profiles. Nucleic Acids Research. 2004 jan;32(90001):91D–94.

[49] Rosenbauer F, Tenen DG. Transcription factors in myeloid development: balancing differentiation with transformation. Nature Reviews Immunology. 2007 feb;7(2):105–117.

[50] Somasundaram R, Prasad MAJ, UngerbÃ¤ck J, Sigvardsson M. Transcription factor networks in B-cell differentiation link development to acute lymphoid leukemia. Blood. 2015 jul;126(2):144–152.

[51] Orkin SH. Transcription Factors and Hematopoietic Development. Journal of Biological Chemistry. 1995 mar;270(10):4955–4958.

[52] Colaprico A, Silva TC, Olsen C, Garofano L, Cava C, Garolini D, et al. TCGAbiolinks: an R/Bioconductor package for integrative analysis of TCGA data. Nucleic Acids Research. 2015 dec;44(8):e71–e71.

[53] Weirauch MT, Yang A, Albu M, Cote AG, Montenegro-Montero A, Drewe P, et al. Determination and Inference of Eukaryotic Transcription Factor Sequence Specificity. Cell. 2014 sep;158(6):1431–1443.

[54] Schep A. motifmatchr: Fast Motif Matching in R; 2018. R package version 1.4.0.

[55] Eferl R, Wagner EF. AP-1: a double-edged sword in tumorigenesis. Nature Reviews Cancer. 2003 nov;3(11):859–868.

[56] Bioconductor Package Maintainer <Maintainer@Bioconductor Org>. ExperimentHub. Bioconductor; 2017.

[57] Marcel Ramos LW. curatedTCGAData: Curated Data From The Cancer Genome Atlas (TCGA) as MultiAssayExperiment Objects. Bioconductor; 2017.

[58] R Core Team. R: A Language and Environment for Statistical Computing. Vienna, Austria; 2018. Available from: https://www.R-project.org/.

[59] Kassambara A, Kosinski M, Biecek P. survminer: Drawing Survival Curves using ‘ggplot2’; 2019. R package version 0.4.6. Available from: https://CRAN.R-project.org/package=survminer.

[60] Grambsch PM, Therneau TM. Proportional hazards tests and diagnostics based on weighted residuals. Biometrika. 1994;81(3):515–526.

[61] Therneau TM. A Package for Survival Analysis in S; 2015. Version 2.38. Available from: https://CRAN.R-project.org/package=survival.

[62] Terry M Therneau, Patricia M Grambsch. Modeling Survival Data: Extending the Cox Model. New York: Springer; 2000.

[63] Holm S. A simple sequentially rejective multiple test procedure. Scandinavian Journal of Statistics. 1979;6(2):65–70.

[64] Ma S, Ogino S, Parsana P, Nishihara R, Qian Z, Shen J, et al. Continuity of transcriptomes among colorectal cancer subtypes based on meta-analysis. Genome Biology. 2018 sep;19(1).

[65] Chikina MD, Troyanskaya OG. An effective statistical evaluation of ChIPseq dataset similarity. Bioinformatics. 2012 jan;28(5):607–613.

[66] Breeze CE, Reynolds AP, van Dongen J, Dunham I, Lazar J, Neph S, et al. eFORGE v2.0: updated analysis of cell type-specific signal in epigenomic data. Bioinformatics. 2019 jun;.

[67] Yu G, Wang LG, He QY. ChIPseeker: an R/Bioconductor package for ChIP peak annotation, comparison and visualization. Bioinformatics. 2015 mar;31(14):2382–2383.

[68] Wang Z, Civelek M, Miller CL, Sheffield NC, Guertin MJ, Zang C. BART: a transcription factor prediction tool with query gene sets or epigenomic profiles. Bioinformatics. 2018 mar;34(16):2867–2869.

[69] Corces MR, Buenrostro JD, Wu B, Greenside PG, Chan SM, Koenig JL, et al. Lineage-specific and single-cell chromatin accessibility charts human hematopoiesis and leukemia evolution. Nature Genetics. 2016 aug;48(10):1193–1203.

[70] Hansen KD, Irizarry RA, Wu Z. Removing technical variability in RNA-seq data using conditional quantile normalization. Biostatistics. 2012 jan;13(2):204–216.

[71] Hinz S, Magheli A, Weikert S, Schulze W, Krause H, Schrader M, et al. Deregulation of EZH2 expression in human spermatogenic disorders and testicular germ cell tumors. World Journal of Urology. 2009 dec;28(5):631–635.

[72] Singh R, Fazal Z, Corbet AK, Bikorimana E, Rodríguez JC, Khan EM, et al. Epigenetic Remodeling through Downregulation of Polycomb Repressive Complex 2 Mediates Chemotherapy Resistance in Testicular Germ Cell Tumors. Cancers. 2019 jun;11(6):796.

[73] Suva ML, Riggi N, Janiszewska M, Radovanovic I, Provero P, Stehle JC, et al. EZH2 Is Essential for Glioblastoma Cancer Stem Cell Maintenance. Cancer Research. 2009 nov;69(24):9211–9218.

[74] Cheng T, Xu Y. Effects of Enhancer of Zeste Homolog 2 (EZH2) Expression on Brain Glioma Cell Proliferation and Tumorigenesis. Medical Science Monitor. 2018 oct;24:7249–7255.

[75] Farlik M, Halbritter F, MÃ¼ller F, Choudry FA, Ebert P, Klughammer J, et al. DNA Methylation Dynamics of Human Hematopoietic Stem Cell Differentiation. Cell Stem Cell. 2016 dec;19(6):808–822.

[76] Gomez L, Odom GJ, Young JI, Martin ER, Liu L, Chen X, et al. coMethDMR: accurate identification of comethylated and differentially methylated regions in epigenome-wide association studies with continuous phenotypes. Nucleic Acids Research. 2019 jul;47(17):e98–e98.

[77] He H, Sinha I, Fan R, Haldosen LA, Yan F, Zhao C, et al. c-Jun/AP-1 overexpression reprograms ER signaling related to tamoxifen response in ER-positive breast cancer. Oncogene. 2018 feb;37(19):2586–2600.

[78] Miranda TB, Voss TC, Sung MH, Baek S, John S, Hawkins M, et al. Reprogramming the Chromatin Landscape: Interplay of the Estrogen and Glucocorticoid Receptors at the Genomic Level. Cancer Research. 2013 jun;73(16):5130–5139.

[79] Jaffe AE, Murakami P, Lee H, Leek JT, Fallin MD, Feinberg AP, et al. Bump hunting to identify differentially methylated regions in epigenetic epidemiology studies. International Journal of Epidemiology. 2012 feb;41(1):200–209.

